# PlantServation: time-series phenotyping using machine learning revealed seasonal pigment fluctuation in diploid and polyploid *Arabidopsis*

**DOI:** 10.1101/2022.11.21.517294

**Authors:** Reiko Akiyama, Takao Goto, Toshiaki Tameshige, Jiro Sugisaka, Ken Kuroki, Jianqiang Sun, Junichi Akita, Masaomi Hatakeyama, Hiroshi Kudoh, Tanaka Kenta, Aya Tonouchi, Yuki Shimahara, Jun Sese, Natsumaro Kutsuna, Rie Shimizu-Inatsugi, Kentaro K. Shimizu

## Abstract

Long-term field monitoring of leaf pigment content is informative for understanding plant responses to environments distinct from regulated chambers, but is impractical by conventional destructive measurements. We developed PlantServation, a method incorporating robust image-acquisition hardware and deep learning-based software to analyze field images, where the plant shape, color, and background vary over months. We estimated the anthocyanin contents of small individuals of four *Arabidopsis* species using color information and verified the results experimentally. We obtained >4 million plant images over three field seasons to study anthocyanin fluctuations. We found significant effects of past radiation, coldness, and precipitation on the anthocyanin content in the field. The synthetic allopolyploid *A. kamchatica* recapitulated the fluctuations of natural polyploids by integrating diploid responses. The data support a long-standing hypothesis stating that allopolyploids can inherit and combine the traits of progenitors. PlantServation pipeline facilitates the study of plant responses to complex environments termed “*in natura*.”

## Introductionx

Plants in complex environments (hereafter referred to as “*in natura*”) are exposed to complex environments with multiple abiotic and biotic factors^1^. Therefore, knowledge from indoor studies, mostly focusing on a single factor, is not necessarily directly transferrable to the field^2–4^. Time-series data from the field are informative for understanding how plants thrive. However, collecting time-series data on plant growth and response in fluctuating environments is labor-intensive and often destructive^5^. Image analysis serves as a non-destructive alternative for time-series data collection in the field; however, the acquisition and analysis of high-resolution time-series images from the field is challenging for several reasons. First, a setup is exposed for months to various weather conditions, such as sunlight, rain, snow, or storms. Such robust systems are often costly^5^. Second, even with fixed-point image acquisition, the camera positions change owing to extreme weather conditions, maintenance work, etc., resulting in the inconsistency of the position of target plants in images from different time points. Third, plant segmentation in acquired images may not be straightforward^6,7^. Plant size, morphology, and color can change over time or are affected by conditions such as snowfall. External conditions such as light intensity, soil texture, and wind disturbance also vary among sites, introducing variations among images from different sites. Fourth, the ideal image resolution to observe a small single individual, such as seedlings of model *Arabidopsis* species, may be difficult to achieve with the currently available cost-efficient methodologies. The resolution of close-up images by drones is typically of the order of centimeters or at best millimeters because of the disturbance of plants caused by the wind that the drones generate^5,8^. In contrast to drones, ground-based systems can easily capture close-up images; however, commercially available products are intended for large-scale fields and are not cost-efficient for small-scale research^9^. Yang et al. (2022) and Hawkesford and Lorence (2017) emphasized that it is important to decrease the cost of phenotyping to promote further research^5,10^. Finally, the amount of collected data can be large, resulting in a long processing time for analysis. Overcoming these challenges and analyzing time-series images of different species in different environments further our understanding of the growth and environmental responses of plants.

Deep neural network (DNN) is a powerful tool for analyzing complex images. DNN is increasingly being deployed to process large image datasets in diverse disciplines, from medical science to engineering^11^. In plant science, it has been successfully implemented to segment the target plant or the position of the plant in an image under controlled conditions, where the plant appearance and background are relatively uniform^12,13^. The analysis methods established for plants with relatively simple shapes under controlled conditions are not directly applicable to complicated images from the field. Large variations in the target plant, light, and background need to be covered in annotating the images from the field to prepare a training dataset for DNN, which can be laborious when performed manually. In addition, the best DNN architecture depends on variations in the images. Various DNN architectures are available with different strengths. For example, U-Net has been used for the segmentation of plants from the top view^6,13,14^ as well as roots in soil^15^. Other architectures that are successful in other disciplines can also be promising for plant image analysis, e.g., SINet in segmenting camouflaged animals^16^ or DANet in detecting fine structures such as human veins^17^. Thus, to efficiently analyze complex images from a field within a realistic workload, it is necessary to reduce the manual annotation effort and select the DNN architecture whose strength best suits the analysis of the features in the target images^6,18^. The application of DNN to high-resolution image analysis of plants in the field enables the identification of diverse biological questions, including evolution, ecology, and environments.

Accumulation of anthocyanin pigment is induced by various environmental conditions and is considered a stress marker^19–21^. Laboratory studies of the model plant *Arabidopsis thaliana* have shown that anthocyanin in leaves increases in response to various external stresses, such as intense light, cold temperature, and drought, as a protection against oxidation, making the plant appear reddish^22,23^. In contrast, even in *A. thaliana*, little is known about the mechanism of anthocyanin accumulation in complex field environments in which air temperature, radiation, and precipitation fluctuate^24^. Furthermore, widespread variation of anthocyanin content within and among species suggests its evolutionary significance in adaptation and speciation^19^.

Allopolyploid speciation occurs through hybridization between different species with genome duplication. Its prevalence among natural and crop plant species has stimulated discussions and debates regarding the advantages and disadvantages of allopolyploid species^25–27^. Since the end of the 20th century, a major focus of the polyploid study has been on genome-wide mutations that are induced at the time of polyploidization termed “genome shock”. However, recent reports have shown a lack of genome shock in *Arabidopsis* and grass polyploids^28,29^, suggesting that it is not essential for polyploid adaptation. Instead of novel mutations, environmental responses of diploid progenitor species can be inherited and combined in allopolyploid species that have been originally discussed in plant evolutionary and systematics studies^27,30–32^. Soltis et al. (2016) have emphasized that the paucity of model polyploid species that integrate functional and ecological data is a major barrier to testing evolutionary and ecological hypotheses on polyploidy^27,32^. In the model genus *Arabidopsis*, the allotetraploid species *A. kamchatica* is emerging as a model polyploid species, which was derived from two diploid progenitors, *A. halleri* and *A. lyrata*^33^. In addition to natural *A. kamchatica* genotypes, synthetic *A. kamchatica* plants can be used to examine the effects of environmental responses inherited from progenitors^34^. The natural distribution range of *A. kamchatica* is wider than that of diploid progenitors, both in latitude and altitudes^35,36^. Physiological and transcriptome experiments in regulated laboratory conditions showed that *A. kamchatica* inherited the gene expression pattern associated with the cold response from the diploid progenitor *A. lyrata* that was distributed in colder habitats than the other progenitors^37,38^. From the diploid progenitor *A. halleri*, *A. kamchatica* inherited the gene expression patterns responsible for zinc hyperaccumulation and tolerance^39^. Zinc concentration analysis of soils from natural habitats showed that *A. kamchatica* can tolerate moderately contaminated soil, suggesting that the allopolyploid inherited adaptive environmental tolerance of *A. halleri* in natural fields^40^. In contrast to relatively stable natural environments such as soil metal concentrations, time-series field observations are critical for capturing plant reactions to fluctuating meteorological conditions.

In this study, we established PlantServation, a method for image acquisition and analysis that consists of hardware and software. Using the robust yet inexpensive image acquisition system with an RGB camera, we collected daily images of four *Arabidopsis* species grown in the field for five months each for three years. We developed an efficient image analysis pipeline using DNN by registering the position of individual plants in time-series images by augmenting annotation data and comparing the performance of multiple DNN architectures. We estimated the time-series anthocyanin content using leaf color information from PlantServation and verified them empirically. To demonstrate the power of the time-series pigment data obtained by PlantServation, we addressed two biological questions: (1) How do air temperature, radiation, and precipitation affect the anthocyanin content in *Arabidopsis* species in complex field environments? (2) Does the synthetic polyploid *A. kamchatica* recapitulate the seasonal fluctuation of anthocyanin in natural polyploids, and how is it associated with those of diploid progenitors?

## Results

### Image acquisition in field

We established an inexpensive image acquisition system that endured in the field for five months during the growing season over three years. The hardware part of PlantServation was set up in common gardens in Switzerland and Japan using commercial polytunnel skeletons and weather-resistant RGB cameras (RICOH WG-40) positioned 150 cm from the ground to collect the top-view images (Fig. 1a). Normal camera batteries do not last longer than a few weeks in our environment. For a stable power supply and to save the labor of exchanging batteries frequently, we replaced camera batteries with custom-made direct current (DC) couplers that could be connected to a common power source after conversion to alternating current (AC) (Fig. 1a, b). A commercial uninterruptable power supply (UPS) provides emergency power in unexpected power breaks (Fig. 1a). The use of a flat cable enabled a connection between the DC coupler and cable while closing and sealing the camera lid (Figs. 1a, S1). The support bars were fixed to each other and to the polytunnel skeletons using moving scaffolding clamps, whereas the DC cables were fixed to the support bars using cable ties (Fig. 1c). To minimize the misalignment of cameras, we inserted a rubber sheet between the camera holder and support bar. We installed a custom-made lid to protect the cameras from snow, rain, and radiation (Fig. 1d). To avoid herbivores, birds, and pollinators, the polytunnel skeletons were covered with mesh sheets, with the top part kept open during winter in Switzerland to allow snowfall and to prevent damage to the polytunnel skeletons by seasonal winds (Fig. 1e). Altogether, the expense for the hardware part of PlantServation with eight cameras to observe 384 plants was ca. USD 2,600.

**Fig. 1.**
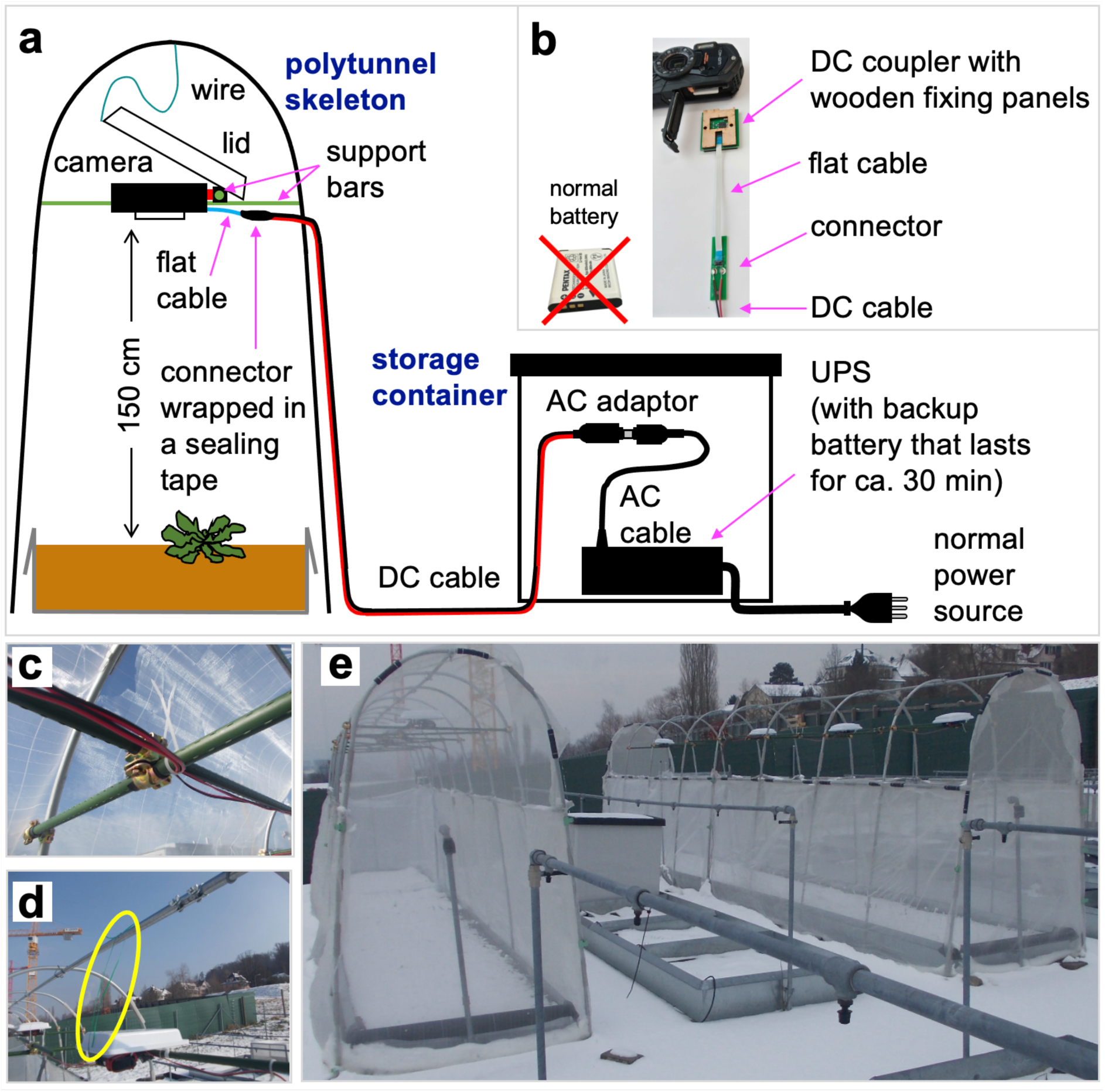
The set-up for image acquisition. **a** Illustration of the image acquisition system with constant power supply. A wire connected with a lid was stretched out when the lid was lowered fully. Instead of a normal battery, a custom-made DC coupler was placed in a RICOH WG40 camera using a flat cable. A UPS was installed as a temporal power source in case of power cut. **b** Close-up of a DC coupler with wooden fixing panels (one for each side of the DC coupler), a flat cable, a connector, and a DC cable. A DC coupler was used instead of a normal battery. A wooden fixing panel and a DC coupler were inserted together in the camera battery slot to position the DC coupler stably. Details of the set-up are shown in Fig. S1. **c** Support bars with DC cables fixed using cable ties and a movable scaffolding clamp with black rubber sheet in between. **d** A wire (yellow oval) lifting the camera lid to avoid contact with camera. **e** Overview of the image acquisition system. The polytunnel frames were covered with mesh, with top mesh being removed during winter in Switzerland as seen in the photo.

Using PlantServation, we obtained approximately 4032,000 images of target plants (12 genotypes × 20 replicates × 2 sites × 16–24 images/day × 150 days/year × 3 years). These were captured using five cameras at each of the Swiss and Japanese sites. Each image had 16M (4608 × 3456) pixels with a pixel range of 0 to 255 in the 8-bit sRGB color space and 1 pixel corresponding to approximately 0.45 mm and included the top view of the 48 target plants as one plot (Fig. 2a). The 48 target plants consisted of four blocks, each of which consisted of 12 genotypes representing four species of *Arabidopsis* (Table S1): the model species *A. thaliana*, natural and synthetic allotetraploid *A. kamchatica*, and its diploid progenitors *A. halleri* and *A. lyrata*^33,36, 41–43^. The newly synthesized *A. kamchatica*, along with its progenitor genotypes, enabled the comparison of polyploids shortly after emergence with those long after establishment and generations under natural selection, while the inclusion of the model plant *A. thaliana* facilitated the interpretation of results in light of previous molecular and physiological studies^38^.

**Fig. 2.**
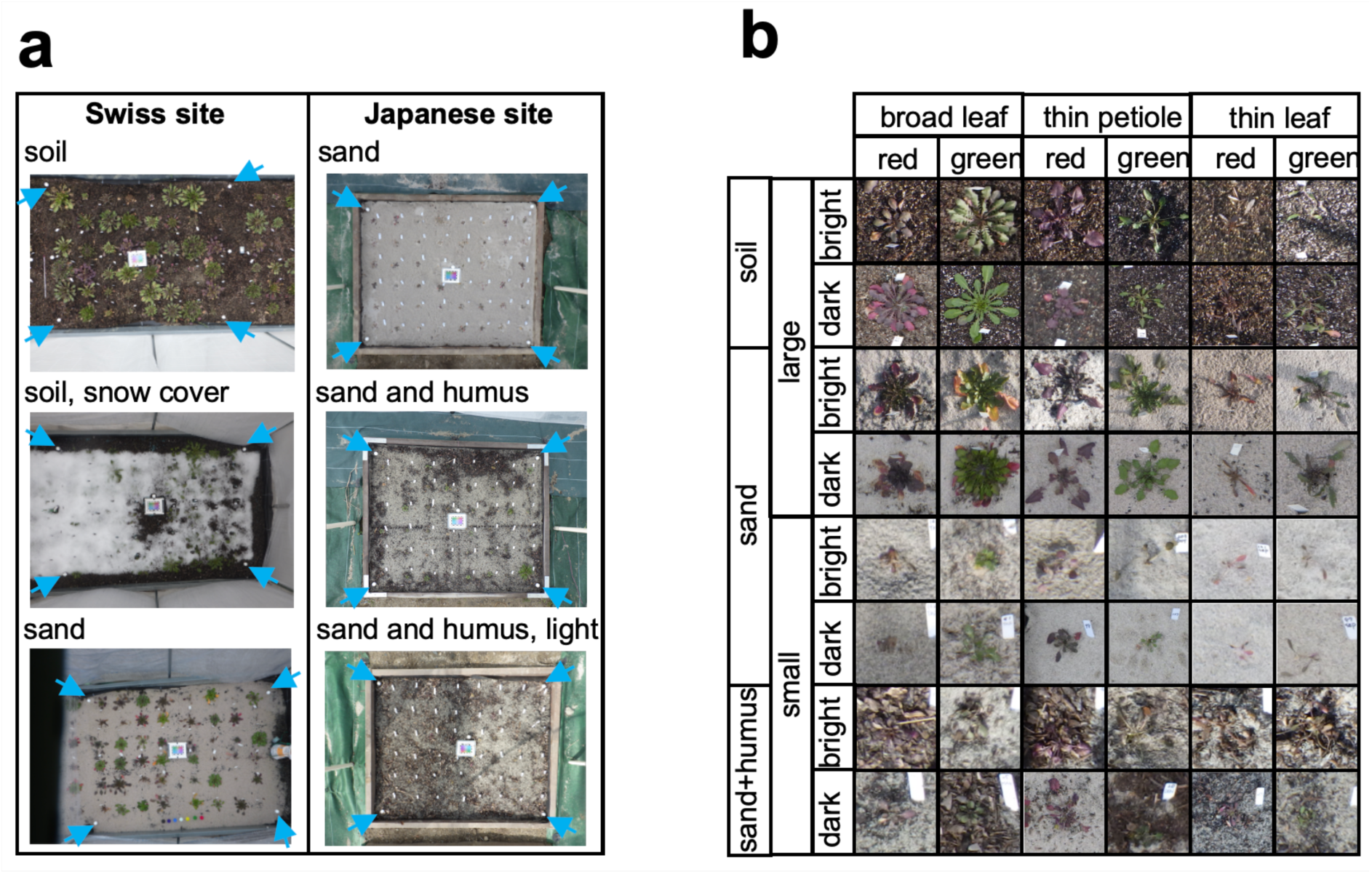
Examples of images of *Arabidopsis* collected in field. **a** Original image from the Swiss site (left column), and from the Japanese site (right column). Background (soil, sand, humus), surroundings (mesh around the plot, frames around the plot, or sheet on the ground), and the amount of noise (e.g., snow seen in the middle images from the Swiss site, strong light on the right half of the bottom images from the Japanese site) varied among images. Blue arrows indicate white marbles placed at the corners of the target area. **b** Close-up images representing the diversity in light condition, in background, and in color, shape, and size of plants

The acquired images contained large variations in layout, background, and target plants. The layout of the frame around the plots varied between the Swiss and Japanese sites, whereas the background (soil, sand, and humus) varied between and within sites (Fig. 2a). Variation also existed at the target plant level with respect to light conditions, background, color, shape, and size of plants (Fig. 2b). In both sites, four white marble balls of 2.5 cm diameter, fixed to wooden or metal nail inserted in the ground, were placed at the corners of the rectangular area containing the target plants (Fig. 2a). These were used as markers for adjusting the position in the images to be consistent throughout the time points (see Step 2 in **Image analysis pipeline** for an overview and **Methods** for details). Using our economical image acquisition system, we successfully collected daily images of different genotypes of *Arabidopsis* in the field over several months.

### Image analysis pipeline

In the PlantServation software, images from each camera were processed using a pipeline implemented in Python (Fig. 3a). Here, we provide a brief overview of our pipeline and describe the details in the **Methods** section. In Step 1 (Fig. 3a), we selected up to four images per day by thresholding the pixel values and setting a time window close to midday to reduce the variation in brightness among the images, yielding ca. a total of 740,432 individual plant images.

**Fig. 3.**
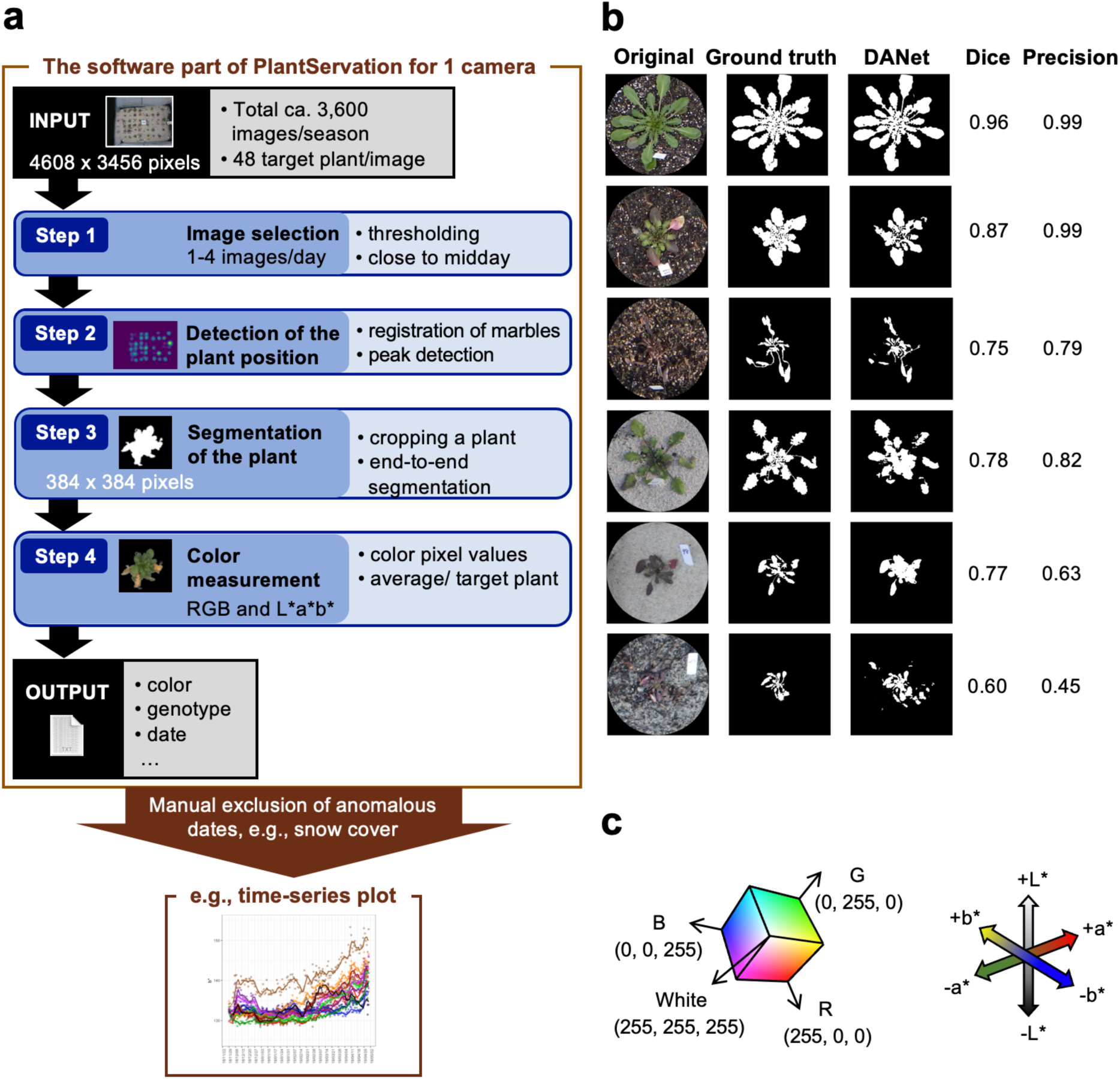
Image analysis pipeline and its performance. **a** The workflow of the image analysis. 16-24 images per camera were taken per day for ca. 150 days per season. Five cameras were set up in two sites for three seasons. **b** The evaluation result of the outcome of the machine learning using DANet. Images varying in plant size, shape, and color and background were examined concerning Dice and Precision. Six representative images are shown. For each plant, the original image with circular mask, ground truth, DANet segmentation outcome, and the score for Dice and Precision are shown. From top to bottom: a green plant with soil background in the Swiss site, a plant with green and dark leaves with soil background in the Swiss site, a dark plant with soil background in the Swiss site, a green plant with sand background in the Swiss site, a dark plant with sand background in the Swiss site, a dark plant with humus background in the Japanese site. **c** RGB color space (left) and L*a*b* color space (right).

In the field, the cameras inevitably move owing to wind and during maintenance, resulting in inconsistencies in the position of the target plants among the images. To address this issue, in Step 2, we performed registration and compilation of the images from different time points and defined the plant position as the peak where the center of the plant was most frequently detected (Fig. S2).

Once the center of the plant was defined, we segmented the plants in Step 3 after cropping individual plants from the image using end-to-end segmentation with DANet that was the best among the five DNN architectures examined (Fig. S3, Table S2). We used a custom-made training dataset consisting of 7,500 images augmented from 225 manually labeled images (Fig. S4).

We examined the performance of our pipeline using DANet by analyzing 33 plant images that were not used to build the pipeline and by comparing the outcome with the ground truth, the plant area marked by humans. The pipeline worked reasonably well in segmenting different background types at the Swiss (soil or sand) and Japanese (sand) sites (Table S3). The performance of the segmentation was slightly higher for the soil background (Fig. 3b). When the color of the plant and that of the background were similar, the Dice coefficient was lower; however, this did not hinder the subsequent color analysis, the major purpose of this study (Fig. 3b).

In Step 4, we color-converted the pixel values in RGB of the segmented plant area to those in L*a*b* (Fig. 3c). We subsequently calculated the average value of L*a*b* per target plant. **Supplementary Movies 1**–**3** show examples of time-series compilation of a segmented plant area.

Finally, the pipeline outputs color information, genotype, and date for each target plant. We excluded anomalous data from further analyses by referring to the field record, for example, on snow cover and plant death, by a Z-score threshold calculated with nearest neighbor interpolation and by visually examining the original image for individuals whose time-series plots deviated from the norm (Fig. S5; see **Methods** for further details).

### Experimental validation of color-based estimation of anthocyanin content

To estimate the anthocyanin content from the color information of images, we used another set of plants to collect color information from images and compared it with the actual pigment content measured experimentally. We regressed the relative anthocyanin content per leaf area on L*, a*, and b* using a random forest model that was previously found to be effective in pigment estimation from color in *Arabidopsis* in the laboratory^44^ (Table S4). Among the three color features, a* contributed the most to the variation in anthocyanin, followed by b* and L* (importance: a* 56.4%, b* 22.4%, L* 21.2%). Fitting results indicated that although the anthocyanin content tended to be overestimated in the small value range and underestimated in the large value range, there was a high correlation between measured and estimated anthocyanin content when genotypes were separated (Pearson correlation coefficient *r* > 0.900 for the majority of genotypes, Fig. S6) and pooled (Pearson correlation coefficient, *r* = 0.911, *p* < 0.081, Fig. 4).

**Fig. 4.**
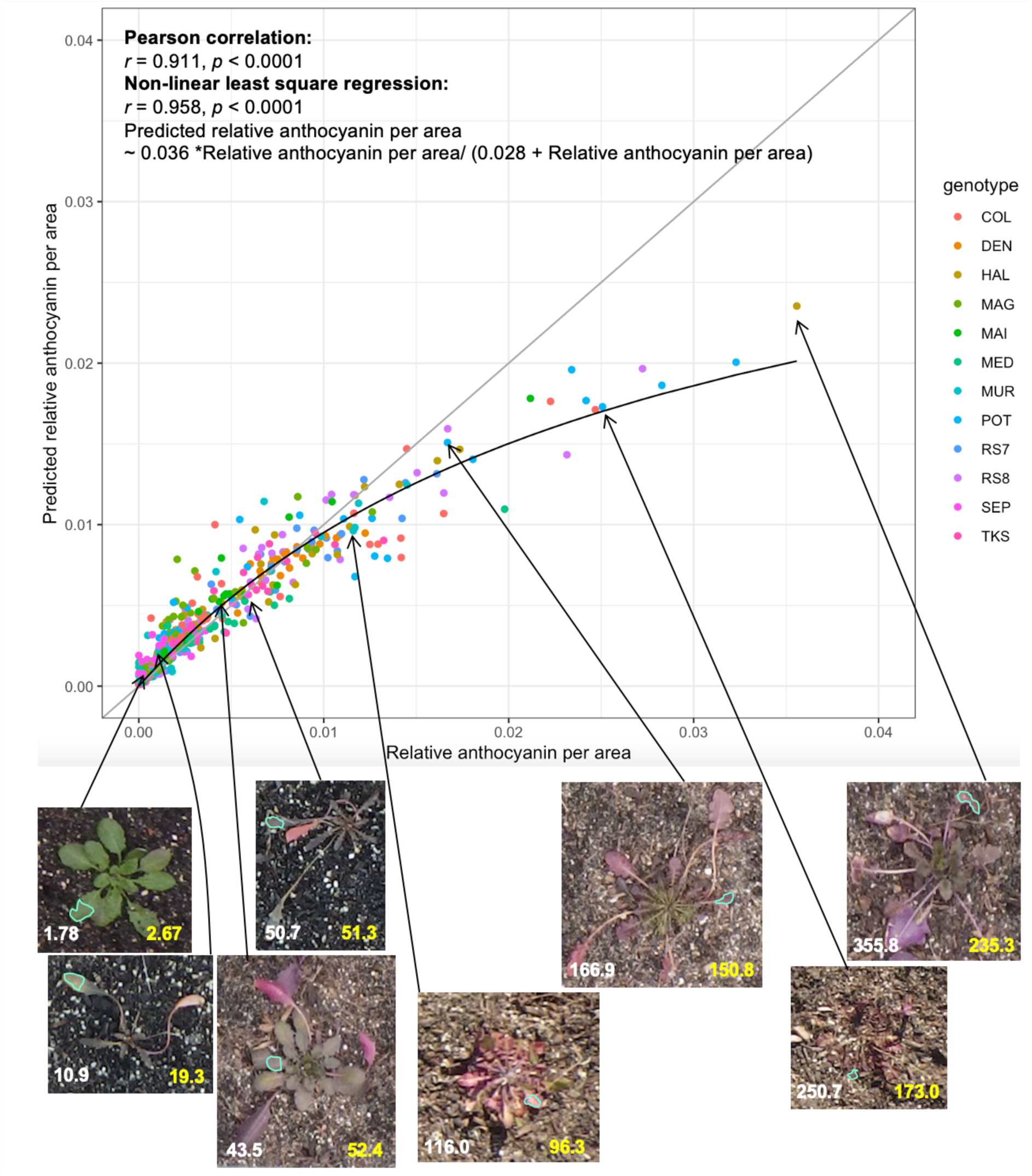
Scatter plot of relative anthocyanin content per area and predicted relative anthocyanin content per area using random forest model. The grey line indicates y = x, while the black line indicates a non-linear least square regression. Images below the plot show representative examples of the estimation of the anthocyanin content using a random forest model. White and yellow numbers indicate measured and estimated values, respectively, where E-4 is omitted for better visibility: e.g., 1.78 = 1.78 E-4. Leaf area subject to the analysis is surrounded in cyan. In the time-series image analysis, whole plant area was analysed using the random forest model built here. *n* = 451.

### Seasonal fluctuation of plant traits

We applied a random forest regression model in the previous section to the time-series images and estimated the anthocyanin content from L*a*b* in the image per time point per genotype per site. To examine the pattern of fluctuation in the estimated anthocyanin content and other traits in the field and the variation among genotypes, we generated time-series plots for each trait per site per genotype (Figs. 5, S7–9). The values were averaged among images when there were multiple images per day.

**Fig. 5.**
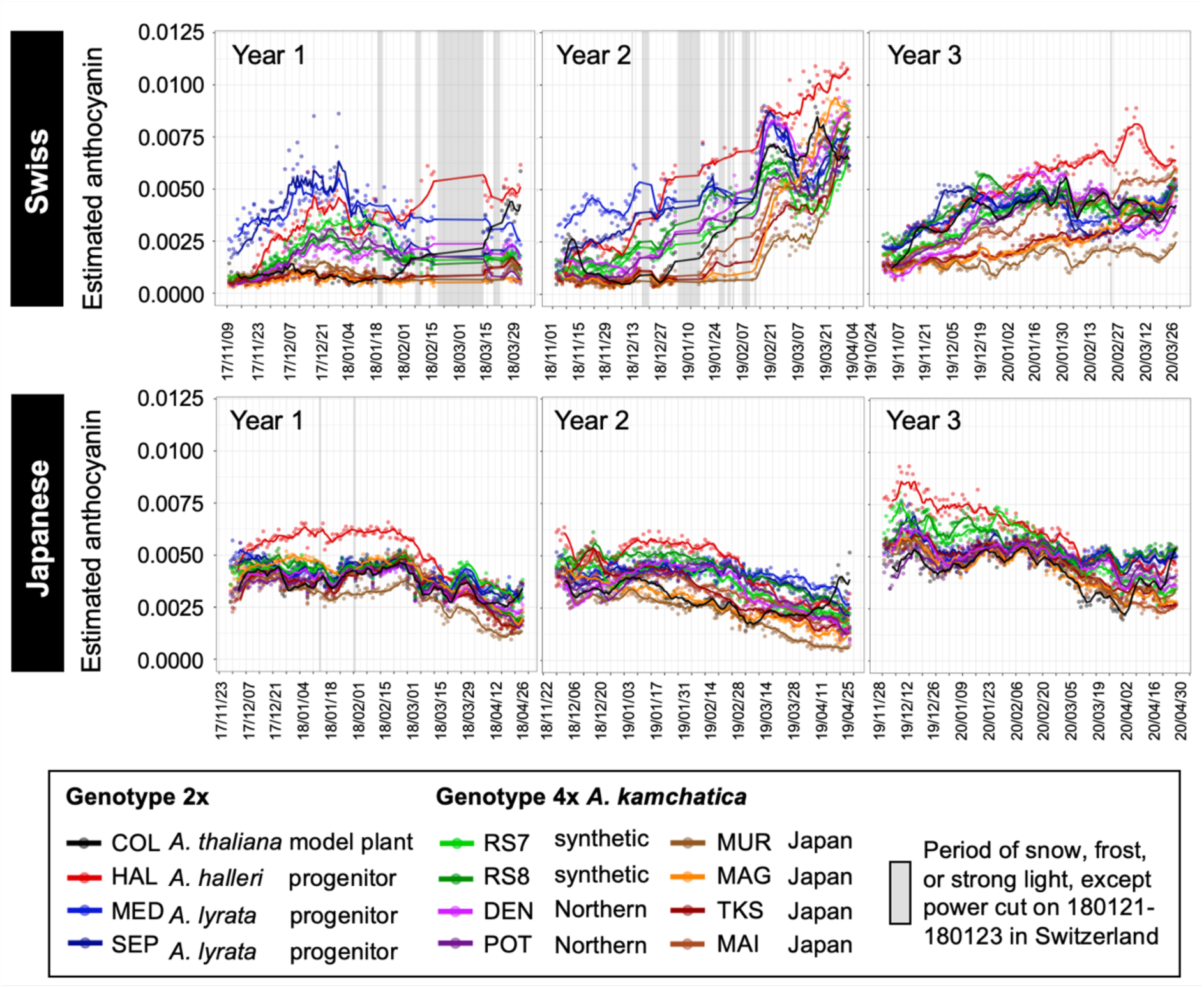
Time-series plots of 5-day moving average of estimated relative anthocyanin content per mm^2^ in plants of 12 *Arabidopsis* genotypes at the Swiss and Japanese sites in three seasons.

The variation in the estimated anthocyanin content throughout the season and among genotypes was more pronounced at the Swiss site than at the Japanese site (Fig. 5). At the Swiss site, the anthocyanin content of most genotypes increased from autumn to winter. The difference between the two sites is consistent with the mild winter conditions at the Japanese site, in which snow rarely falls. These data will be used for analyses in the following two sections.

The a* trend resembled that of the estimated anthocyanin content (Figs. 5 and S7). In contrast to a* and the estimated anthocyanin content, b* and L* showed a reverse pattern over time and in the order of the genotypes according to the value of the variable (Figs. 5, S7–S9). These results were consistent with the large contribution of a* to the estimated anthocyanin content (see the previous section).

### Effect of radiation, coldness, and precipitation on anthocyanin content in *Arabidopsis*

Experiments in regulated chamber conditions showed that anthocyanin in *A. thaliana* is induced by stress treatment at low temperature, strong light, and drought ^22,23^. Using the time-series anthocyanin content estimated in the field, we examined whether environmental conditions had a significant effect on the anthocyanin content of *A. thaliana* and its relatives in the complex natural environments (Fig. 6a). We fitted linear regression models with the estimated anthocyanin content as the response variable and radiation, coldness, and precipitation as explanatory variables. Considering the response time and threshold of plants to environmental cues, we adopted the best parameter combination for window, lag, and temperature thresholds in the past month, as in previous phenological studies^45^ (Fig. 6a). We found that the estimated anthocyanin content was associated strongly with coldness and radiation and relatively weakly with precipitation in most genotypes (Table S5). This suggests that the three stress conditions studied in the laboratories also exhibited significant effects on the field conditions and confirmed that PlantServation workflow is useful for evaluating the environmental effect on plant traits. In addition, the analysis showed that each of the three environmental factors explained 10–60% of the variation in the estimated anthocyanin content. At the Swiss site, the radiation was particularly influential (Fig. 6b). Time-series plots of the model species *A. thaliana* at the Swiss site with the estimated environmental parameters suggested that the estimated anthocyanin content was well associated with radiation, whereas the relationship between rainfall and coldness was not straightforward (Fig. S10). At the Japanese site, coldness contributed significantly to all the genotypes, although the overall proportion of variations explained by the three factors tended to be lower than those at the Swiss site, possibly because of the mild winter with negligible snow. Considered together, the results suggest that coldness, radiation, and precipitation contribute to the anthocyanin content in the field, with the extent of contribution varying among environmental factors, sites, and genotypes. Roughly, half of the variations in anthocyanin content were not explained by the three environmental factors, suggesting the need for further studies on other factors and their combinatorial effects (see **Discussion**).

**Fig. 6.**
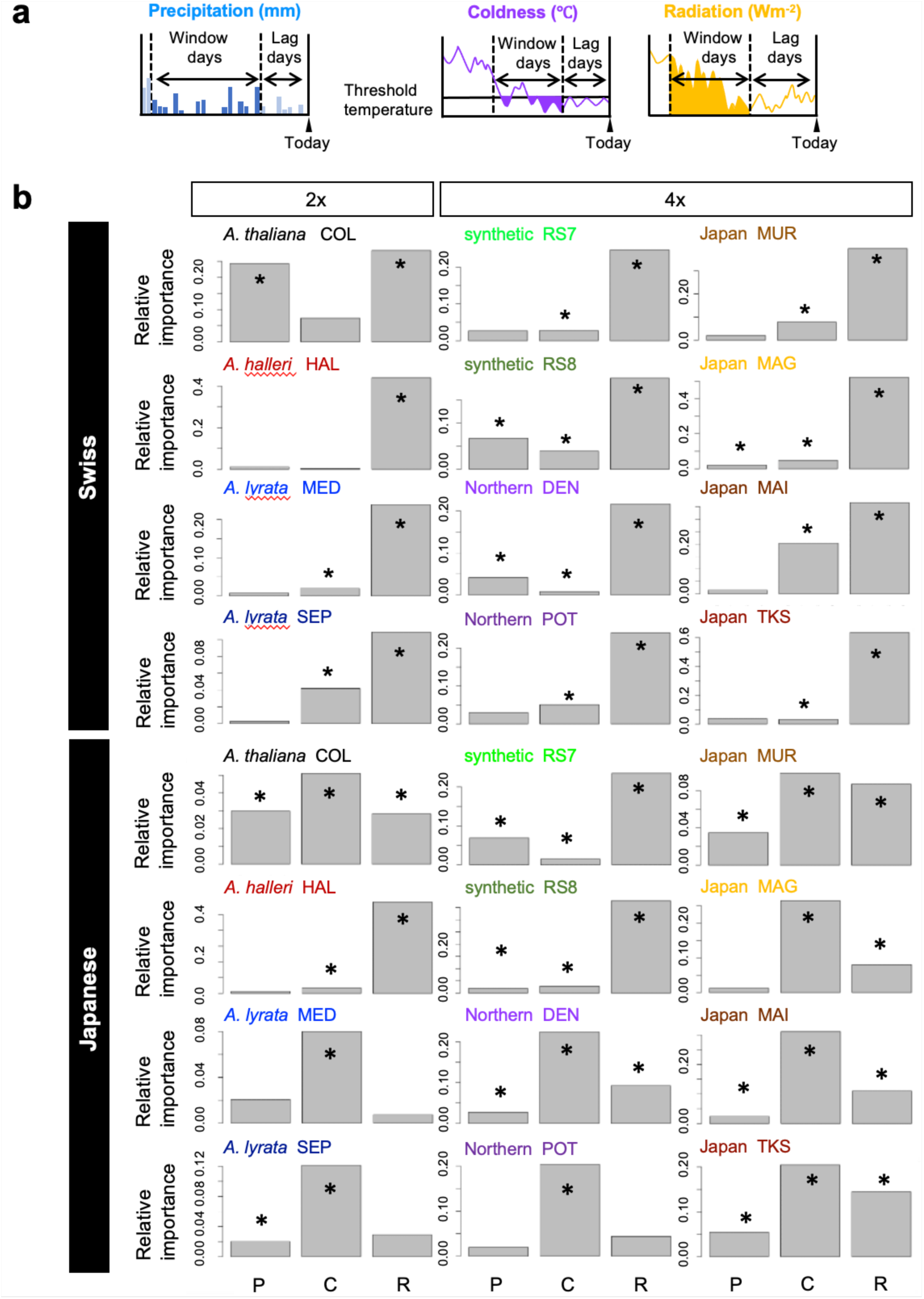
Environmental variables associated with the estimated anthocyanin content. **a** Illustration of the concept of the moving total of precipitation, coldness, and radiation used in the regression analysis. **b** Relative importance of the environmental variables on the estimated anthocyanin content for 12 genotypes of *Arabidopsis* at the Swiss and Japanese sites. P: Precipitation, C: Coldness, R: Radiation. The genotype code in three capital letters are indicated for each genotype. 2x indicates diploid and 4x indicates allotetraploid *A. kamchatica*. Significant variables based on confidence intervals calculated with bias-corrected and accelerated (BCa) bootstrapping are indicated with asterisks. *n* = 663.

### Anthocyanin content in synthetic and natural *Arabidopsis* polyploids

Next, we examined the differences in the estimated anthocyanin content between species and genotypes. We addressed the question of whether synthetic polyploids recapitulate the patterns of natural polyploids and whether they combine the traits of diploid progenitor species. We included two independent synthetic polyploid genotypes of *A. kamchatica* with their diploid progenitors: RS7 derived from HAL (*A. halleri*) and SEP (*A. lyrata*), and RS8 derived from HAL and MED (*A. lyrata*). As natural *A. kamchatica*, we planted four genotypes of Japanese polyploids and two genotypes named Northern polyploid, which is estimated to have originated independently from Japanese polyploids^36^. The Swiss site showed a higher variation among genotypes and seasons, as described above, and they can be grouped roughly into four according to the pattern of fluctuation of the estimated anthocyanin (Fig. 5). First, the diploid progenitors *A. lyrata* (MED and SEP) had a higher content than others in early seasons, followed by intermediate in late seasons. Second, the diploid progenitor *A. halleri* (HAL) showed an opposite trend (intermediate and subsequently higher than the others). Third, the Japanese allopolyploids exhibit low contents throughout the seasons. Fourth, synthetic allopolyploids and natural allopolyploids of northern origin showed an interesting pattern. They did not constitute a simple addition of diploid progenitors but a combination of periods during which they resembled diploid progenitors (Fig. 5). For example, the synthetic allopolyploids showed a trend similar to that of the natural diploid *A. halleri* at the beginning and to the diploid *A. lyrata* later in the second year at the Swiss site. Overall, the trend of the synthetic allopolyploids resembled that of the diploids with a smaller content of the estimated anthocyanin at a given time point.

To compare the trend of the estimated anthocyanin content among genotypes throughout the seasons, we conducted dimension reduction via principal component analysis (PCA) on the estimated anthocyanin for all years from both sites (Fig. 7). The first principal component (PC1) explained 54.3% of the variation in the data. Along PC1, the diploid progenitors were closer to the synthetic allopolyploids, and the synthetic allopolyploids were closer to the natural allopolyploids of northern origin than to the natural allopolyploids from Japan. The second principal component (PC2) explained 15.3% of the variation. Along PC2, two diploid species were located at both tips (Fig. 7).

**Fig. 7.**
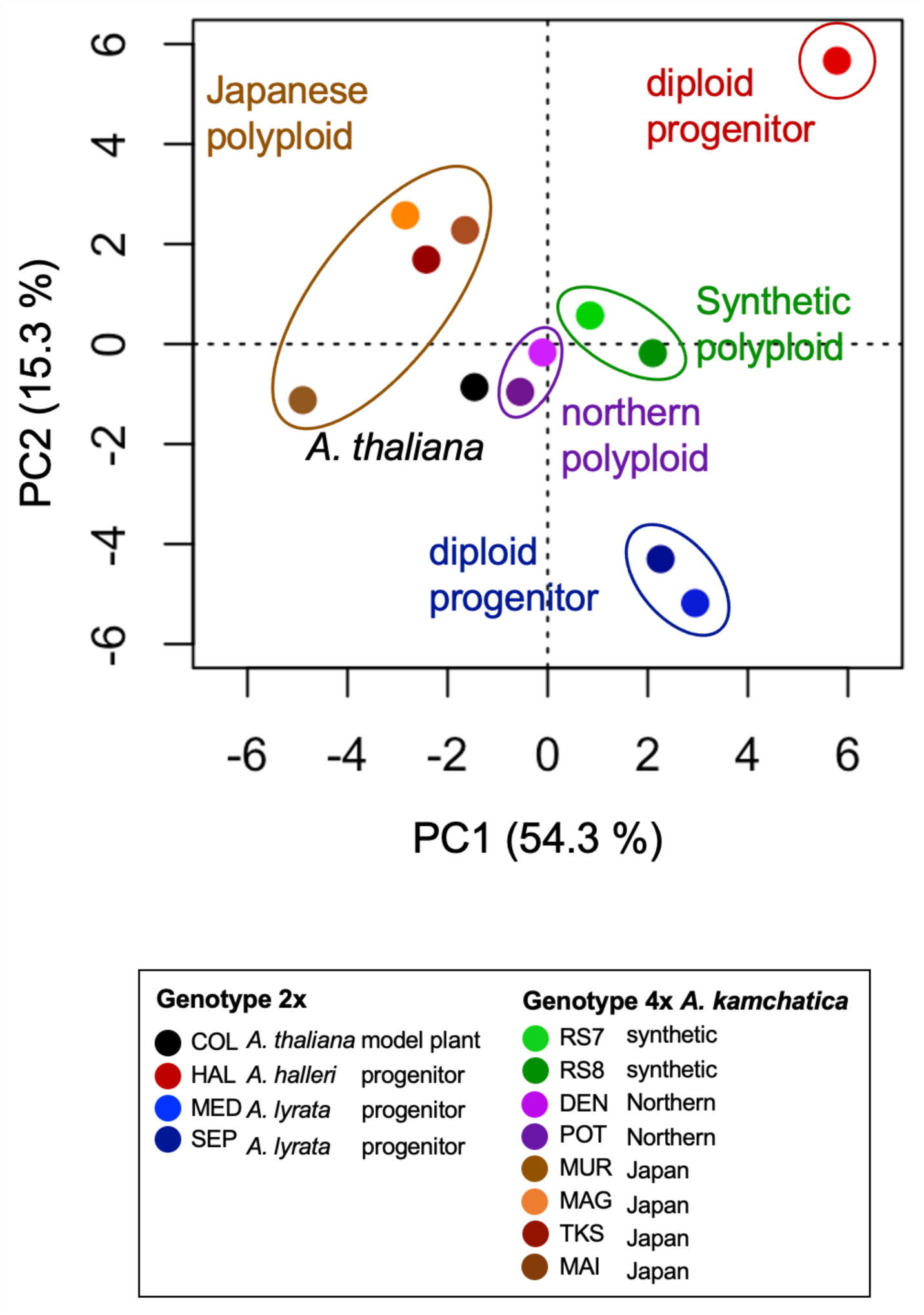
Principal component analysis plot of time-series estimated relative anthocyanin content per mm^2^ for 12 *Arabidopsis* genotypes. The data from the Swiss and Japanese sites in three seasons are merged.

The PCA plot indicated that the two independent synthetic polyploids showed a similar pattern, confirming reproducible changes immediately after polyploidization. Furthermore, the synthetic polyploids were closely related to natural polyploids, that is, two genotypes of northern polyploids (Fig. 7). The other four natural polyploids showed similar PC2 values to synthetic polyploids but diverged at PC1, potentially reflecting their different or older polyploid origin (see **Discussion**). These data suggest that the synthetic polyploids recapitulated the pattern of anthocyanin responses of certain natural polyploids. Furthermore, the synthetic polyploids along with natural northern polyploids were located in the middle of the two diploid progenitor species (Fig. 7), supporting the observations of the inheritance and merger of the traits of the two diploid progenitors. All these patterns persisted when we split the data into two sites, suggesting that they are inherent in the genotypes and are exhibited regardless of the locations of growth (Fig. S11). The grouping of the genotypes in these analyses on genotype averages was generally consistent with that of the analyses at the individual plant level (Fig. S12). The plots of the PC scores with the estimated anthocyanin content for each genotype suggest that some site-by-year combinations, such as the Swiss site Year 1 (yr1) for PC1 and the Swiss and Japanese sites Year 3 (yr3) for PC2, may be particularly influential to the among-genotype difference of the overall PCA plots (Fig. S13).

## Discussion

### Establishment of an analysis method of time-series images from field

Using PlantServation, we successfully distinguished the time-series trends of different plant genotypes and species varying in morphology, color, and size under fluctuating and noisy outdoor conditions via image analysis using DNN. This was achieved by a relatively small manual annotation effort owing to augmentation (Fig. S4). Our data demonstrate that it is possible to quantitatively and non-destructively evaluate plant color at high resolution throughout the season. Hence, we could overcome the challenges in conventional manual data collection and qualitative color evaluation using visual inspection or color charts^46^. Notably, our image acquisition system is robust and simultaneously relatively simple and inexpensive, allowing non-experts to implement the data acquisition without a large investment. Many waterproof, shockproof, freezeproof, and dustproof cameras with interval shooting functionality are available on the market. Any of such tough cameras can be an alternative if a compatible DC coupler can be prepared. All other parts, e.g., support bars, cables, terminals, UPS, of the hardware are also easily available at affordable prices. In this regard, this study paves the way for comparing and classifying the responses of plants of different genotypes and species to seasonally fluctuating environments.

The best DNN architecture for analyzing our dataset was DANet. This could be attributed to the strength of DANet in detecting fine structures such as petioles in our dataset. For our dataset, DANet outperformed SINet, though the latter is widely known for detecting camouflaged objects. Although not optimal for our dataset, U-Net has been shown to perform effectively with small labeling datasets in medical research^47–49^ and has recently been increasingly applied to agriculture for leaf disease symptom diagnostics^50^. Many DNN architectures have been developed in fields other than plant science^11^. When selecting a DNN architecture, widening the search beyond plant science can yield a better solution. Indeed, DANet was originally developed to detect scenes and objects on the street in computer vision and was later successfully applied to detect fine vessels in the human retina in medical research^17,51^. For accurate segmentation, it is important to grasp the critical features of the target image and select a suitable DNN architecture that can detect them, regardless of the type of object to be segmented.

Augmentation achieved reasonable segmentation performance with a relatively small labeling dataset. We used 225 manually labeled images that is markedly less compared with similar indoor studies in which hundreds or thousands of images were labeled^12,13^. Considering the complexity of our images from the field, our approach was highly labor-saving. For similar plant segmentation tasks in future studies, fewer manually labeled data will be sufficient to construct a DNN model using the transfer learning of our learned model as well as the data augmentation strategy^52^. Our data also highlight many difficulties in handling time-series images from the field. Among them, detection of the target plant was the most critical for accurate data acquisition for downstream analysis. In particular, whenever the color and texture of the target plant and background are similar, it is challenging to identify the area of the target plant, even for humans. Even though it might be difficult to avoid such situations, data quality could be improved by addressing other issues. A potential future possibility is to change the segmentation procedure. Segmentation was independently performed for each frame. The incorporation of temporal information into the DNN input may facilitate the segmentation of images that are difficult to segment within a single frame. In this context, the application of Video Object Segmentation methods is potentially promising^52^. Another possibility is the additional implementation of manually obtained information regarding the target plant and its circumstances. Non-target objects in the image, such as snow, can be labeled for training the DNN to distinguish them from the target plant during segmentation (Fig. S5a). Cross-referencing with manual records is effective for issues that are difficult to address by modifying the analysis pipeline. For example, plant death may not be precisely recognized by the DNN when the plant body remains intact (Fig. S5b). In such cases, manual scoring of plant survival can complement the analysis of time-series growth.

Overall, time-series image acquisition in the field inevitably faces a number of predictable and unpredictable challenges, from the similarity in color and texture between target plants and their surroundings to the interference of nontarget objects. Not all the issues would be solvable; however, a priori preventive measures as well as the measures necessary for the image acquisition system should improve the quality of downstream image analysis. An effective understanding of the study system, e.g., the plant life cycle and weather conditions at the study site, would enable selecting the appropriate measures for each case.

### Use of field time-series images to understand seasonal fluctuations in anthocyanin and plant responses to environments

We estimated the time series of leaf anthocyanin content from the color information. As in a previous study on *Arabidopsis thaliana* in the laboratory, a random forest model functioned effectively for our dataset in the estimation of anthocyanin^44^. With the strength of capturing nonlinearity in data, a random forest model can be effective for samples from fields where the environment is heterogeneous, causing noise in the color information. By successfully estimating anthocyanin content from color information, this study opens up the possibility of monitoring plant physiological responses to the environment over months in a non-destructive manner.

Our study demonstrates that time-series data can contribute to the evaluation of plant responses to environments. Regression analyses indicated that in most genotypes, more than one of the cumulative coldness, radiation, and precipitation in the past days to weeks were significantly associated with the estimated anthocyanin content in field. These results are consistent with those of previous studies in the laboratory reporting temperature, light, and drought as key environmental factors affecting anthocyanin accumulation and highlight the importance of considering multiple environmental factors (Fig. 6b)^2,22,23,53^. The linear regression model incorporating lag and window of past temperature suggests that the anthocyanin content is better explained by past temperatures than by the current temperature (Table S5, the extreme would be lag 0 and window 1). Along with gradual changes in anthocyanin content, plants may accumulate past temperature information through transcriptional and epigenetic memory similar to vernalization, seasonal responses, or heat acclimation^1,54–57^. Integration of molecular data with image analysis is valuable for understanding the molecular basis of plant responses *in natura*.

The three environmental factors that were tested based on previous laboratory studies explained roughly half of the variation in the anthocyanin content in fields. Beside noise in the field, the unexplained variations suggest that the linear regression models incorporating the lag and window of the three factors are not adequate for understanding the complex fluctuations in the field. Fine-grained measurement of climatic variables, such as soil moisture, distinct measurement of air and soil temperature, the spectrum of irradiation including UV-irradiation, and the interaction thereof, would explain more variations in anthocyanin content^2,20^. In addition, biotic interactions such as disease and herbivory are known to affect anthocyanin content, although they were not observed in these experiments^20,21^. We suggest that little is still known about plant environmental responses *in natura*. Our pipeline paves the way to search for combinatorial environmental effects along with intrinsic developmental stages and thus, would increase the proportion of anthocyanin changes to be explained.

Time-series monitoring of anthocyanin content in *Arabidopsis* species in the field provided a unique opportunity to address long-standing questions on polyploidy using ecological data^27,32^. First, the fluctuations in anthocyanin content of the synthetic allopolyploid *A. kamchatica* in the field were highly similar to those of certain natural polyploids, that is, northern polyploids. This suggests that synthetic polyploids can recapitulate the polyploid speciation of *A. kamchatica* that is observable in anthocyanin changes. The wide variation among the natural polyploids may be attributed to their local adaptation or independent origins. Molecular population genetic studies suggested that northern polyploids had an independent polyploid origin and possibly originated more recently than Japanese polyploids based on their high similarity to diploid sequences^36,58^. The divergence of Japanese polyploids from synthetic polyploids may be attributed to the longer evolutionary time since polyploidization. In addition, Japanese polyploids may originate from diploid genotypes with different anthocyanin responses.

Second, the analysis supported a long-standing hypothesis stating that the synthetic polyploid can combine the responses of two progenitor species. The two independent synthetic polyploid lines showed similar results, suggesting that the stochastic novel mutation at the polyploidization events (may be called “genome shock”) did not play a significant role^27^. This phenotypic trait analysis is in agreement with transcriptomic studies showing inherited and combined responses of allopolyploid species^27,38,59,60^. Interestingly, the anthocyanin content of the synthetic polyploids tended to be closer to the lower values of the two diploid progenitors. If we can assume that anthocyanin content is a stress indicator, it suggests that synthetic and northern *A. kamchatica* accumulate less anthocyanin than progenitors because they are able to withstand stress throughout the season by obtaining a generalist niche through the combination of progenitors’ environmental responses. Consistent with the observed similar anthocyanin contents of *A. kamchatica* and *A. lyrata* during cold seasons, laboratory studies suggested that *A. kamchatica* inherited cold tolerance and associated gene expression patterns from the progenitor *A. lyrata*^37,38^. To further verify this generalist hypothesis originally proposed by Stebbins, measurements of fitness components are important^30^.

### Potential application of PlantServation

As laboratory settings do not necessarily reproduce field environments, field data are essential for understanding plant responses to complex natural environments^3^. While retaining the simplicity of the image acquisition system reported earlier, PlantServation enables continuous image acquisition in the field over months, capturing changes in leaf color and pigment in seasonally fluctuating environments^61^. We envisage the application of PlantServation in field experiments on model species. The spatial resolution of PlantServation enabled monitoring *Arabidopsis* seedlings at the individual level. Field observations of *A. thaliana* and its relatives will enable the study of plant responses in the field, taking the advantage of a large number of studies in regulated chambers as well as mutant collections. For example, destructive sampling of *A. thaliana* showed that anthocyanin content was reduced in the double mutants of *UVR8* and *CRY1* photoreceptor genes in the field^62^, which can be extended to time-course studies. By connecting to solar power, PlantServation can be deployed at remote natural sites to observe diverse species. Furthermore, PlantServation can be exploited for other studies, including the screening of crops, where trait scoring at an individual plant or a finer level is informative. For example, the drought and iron deficiency responses or disease resistance quantified in previous studies can be monitored for a longer period^63–65^. Furthermore, although it was not the focus of the current study, the combination of our image acquisition system and the image analysis pipeline could also detect differences in morphological features among plants, as observed in the segmentation results (Fig. 3b). Thus, this study should also contribute to fine-scale morphological analyses using field images in future studies.

## Conclusions

We demonstrated that it is possible to estimate the anthocyanin content in plants grown in the field based on the color information obtained from images using an inexpensive photo shooting system and an efficient image analysis pipeline using DNN. Moreover, we showed that time-series field data can provide insights into plant evolution and environmental responses, furthering our understanding of how plants thrive *in natura*.

## Methods

### Study species

We studied 12 genotypes of four species in the genus *Arabidopsis.* The *Arabidopsis thaliana* (2*n* = 10) Col-0 accession was used as the standard experimental strain. *Arabidopsis lyrata* (2*n* = 2*x* = 16) inhabits a circumpolar region, whereas *A. halleri* (2*n* = 2*x* = 16) is distributed in central Europe and the far east^41^. The allotetraploid *A. kamchatica* (2*n* = 4*x* = 32) originated from diploids *A. lyrata* subsp. *petraea* and *A. halleri* subsp. *gemmifera*^36^. *Arabidopsis kamchatica* is widely distributed ranging from Taiwan, Japan, and Siberia in the far east to North America^37^. Two subspecies are recognized in Japan: *A. kamchatica* subsp. *kawasakiana*, which is restricted to sandy shores in the lowlands and *A. kamchatica* subsp. *kamchatica,* which occurs at various altitudes^33,36,42^. Two synthetic *A. kamchatica* samples of laboratory origin were included (Table S1). The genotype RS8 was synthesized by applying colchicine to a seedling of a hybrid of the diploid progenitors *A. halleri* subsp. *gemmifera* (W302 from Japan, called HAL) and *A. lyrata* subsp. *petraea* (named MED after its synonym *Arabis media*); the genome assembly has been reported for both^58,66^. The genotype RS7 was automatically polyploidized from a hybrid of the same *A. halleri* genotype and *A. lyrata* subsp. *Petraea* from another population in Siberia (named SEP after its synonym *Arabis septentrionalis*)^67^.

### Study sites

The study was conducted at the Center for Ecological Research, Kyoto University, Japan (34°58′16″N, 135°57′24″E, 150 m a.s.l.), and at the University of Zurich, Switzerland (47°23′46″N, 8°33′05″E, 150 m a.s.l.).

### Experimental set-up

The experiment was conducted from autumn (seedling stage) to the following spring (end of the vegetative stage), starting in 2017 (yr1), 2018 (yr2), and 2019 (yr3). According to the local phenology of the study species, we transplanted seedlings in Switzerland and Japan in September or October, and November, respectively.

### Plant cultivation

A brief overview of this process is provided below. Additional details are provided in the **Supplementary Note 1**.

#### Swiss site

Seeds were germinated and incubated in a growth chamber (long-day chamber: 22 °C/20 ℃, 16 h:8 h light: dark, RH 60%, light 120–140 μE) or (short-day chamber: 18 °C/16 ℃, 8 h:16 h light: dark, RH 60%, light 120–140 μE, only for *A. thaliana*) for six weeks. We clonally propagated *A. halleri* using small branch segments (1–2 cm) in a long day chamber for five weeks. We acclimated all plants outside under the roof for a week before transplanting them to common garden plots (1 m × 7 m each) filled with Rasenerde (Ökohum GmbH, Switzerland) at the Irchel Campus of the University of Zurich. Only in yr3 did we have a ca. 5 cm layer of quartz sand on the soil surface to have a better color contrast between the plants and ground.

#### Japanese site

The seeds were germinated and incubated in a growth chamber (22 °C/20 ℃, 16 h:8 h light: dark, RH 74%, light 125–145 μE, KOITOTRON HNM-S11) for six weeks or three weeks (only for *A. thaliana*). *A. halleri* was clonally propagated as described above and incubated for five weeks. After one-week acclimation outside under the roof, the seedlings were transplanted into a compartment (W × L × H: 100 cm × 100 cm × 20 cm) filled with a mixture of 4:6 humus (100% Shinshu fall leaves 100% natural fermented products, Koshin Kawara Co. Ltd.) and quartz sand (0.3–0.6 mm). We covered the surface of the soil mixture with a 1–2 cm-thick layer of quartz sand to increase the color contrast between the plants and the ground.

### Experimental design

The plants were placed according to a randomized complete block design with a 15 cm interval to the next plant. Four adjacent blocks (2 × 2) constituted one plot, each of which was monitored using a single camera. The number of plots was five for each of the Japanese and the Swiss sites.

### Image acquisition (PlantServation hardware)

We used a RICOH WG-40 camera resistant to water, shock, dust, and freeze (protection level IP68) with autofocus and no flash modes. The detailed settings of the photo shoot are shown in Table S6. The cameras were fixed onto frame bars using mounting tools (RICOH O-CM1472 and O-CH1470) that were specific to the camera. The acquired images were stored on an SD card and manually downloaded to a PC using a cable connection.

Continuous interval shooting throughout the season (16 images per day in yr1 and yr2, and 24 images per day in yr3) was enabled by the customized power supply system (Fig. 1a). The combined use of an AC adaptor (9 V) and a house-designed DC coupler converted the voltage to 5 V to operate the camera (Fig. 1b, see Fig. S1 legend for detailed design). A wooden fixing panel was placed next to the DC coupler to stabilize the position of the thin DC coupler in the camera battery slot. The flat cable connected to the DC coupler came out from the camera with the lid closed (Figs. 1a, S1) with sealant (3M Gel Coating GC-TCORL) on the rim of the lid to prevent corrosion. Outside the camera, the flat cable was connected to a 10 m-long DC cable. The connecting points (‘connector’ in Fig. 1b) were sealed with a self-fusing tape for sealing and insulation (Fig. 1a). We bound four DC cables, each of which was obtained from one camera, using a power barrel connector jack connected to an AC adaptor. The UPS station supplied power for up to 30 min in the case of a power break to prevent the interruption of photo shooting (Fig. 1a).

### Image analysis (PlantServation software)

Fig. 3 summarizes the workflow of the image analysis conducted using in-house Python scripts (version 3.6.6).

Step 1: We selected up to four images per day that satisfied the following criteria: First, thresholding allowed only images with maximum and average pixel values greater than 80 and 10, respectively, to be analyzed. Second, the time at which an image was acquired was restricted to 10:00–14:00. These filterings reduced the variation in brightness and sunlight direction, and the contamination of accidentally dark images.

Step 2: To adjust the position of the target plants and detect each plant location, we first detected white marbles at the corners of the plot using a DNN (ResNet), registered the image series via homography transformation, matching the marbles to reference the marbles which were determined by averaging the position of each marble, and subsequently downsized each to an image of 1152 × 864 pixels (Fig. S2). Each of these low-resolution images was divided into nine patches of 384 × 384 pixels, and segmentation was applied to roughly assign 1 and 0 to the plant and background regions, respectively. Subsequently, we adopted both areas for overlapping parts (‘AND’) to fit 1152 × 864 pixels. This image was enlarged four times in length (4608 × 3456). This series of resizing and processing steps contributes to shortening the total processing time. Subsequently, we overlaid the mask images (where the object and rest were assigned to 1 and 0, respectively, to generate binary images) acquired at different time points and obtained one summed image with local peaks. Each local peak corresponds to the center of the plant from a given time point. The position of the local peaks was regarded as the position of the target plant in the image. Because the peak was distinctly separated from the background value and exhibited a distinct value, peak detection was straightforwardly performed using the peak_local_max function in the skimage feature library (https://scikit-image.org/docs/dev/api/skimage.feature.html). After peak detection, we examined whether the plants were correctly detected, and when not correctly detected, we manually identified the plant or removed the plant that had been incorrectly identified.

Step 3: Once the plant position was determined, we performed segmentation of the target plants in each 4608 × 3456 pixel image (Fig. 3). Although a patch-based segmentation method using a convolutional neural network was effective for detecting fine features in a previous study^68^, it was time-consuming and missed detailed features of our dataset; therefore, we used an end-to-end segmentation method with DNN using the library pytorch (Fig. S3).

The training dataset for the segmentation of the target plants was prepared as shown in Fig. S4. First, using the annotation tool Labelme (https://github.com/zhong110020/labelme), we labeled 225 images with soil background at the Swiss site consisting of plants with diverse colors, sizes, and morphologies. The 225 labeled images were augmented by rotating, shifting, scaling, and changing the brightness and contrast to yield 4,100 training images. We used the same 225 labeled images to generate 3,400 training images for the sand background at the Swiss and Japanese sites. For this, we overlaid a labeled plant image on a randomly selected background image, from which we cut out the area of the plant. Thereafter, for each of the 7,500 training images, we cut out an area of 384 × 384 pixels containing the target plant and cropped it by cutting out a circular area with a diameter of 384 pixels with the plant position as the center. This cutting procedure was effective in excluding neighboring plants. In later growth stages, some of the neighboring leaves may be included, but they are considered minor for the effect on color estimation in comparison to the fully grown focal plant. This image was used as the input for training using the best DNN architecture for our dataset. The best DNN architecture was identified as the method that yielded the most accurate segmentation results using 4,100 training data points for the soil background in the Swiss site. We compared the standard U-Net^14^, U-Net with pre-trained ResNet-101^69^ or EfficientNet-B7^70^ as a backbone, SINet^16^, and DANet^51^. For all DNN architectures, we split the 7,500 input data into 68%, 12%, and 20% corresponding to training, validation, and test set, respectively. DANet performed best with our dataset and was used in the analysis (Table S2).

After the segmentation of the plant, post-processing was performed to remove noise. First, the output image of the DANet was subjected to Gaussian filtering at sigma = 1 (pixel). After filtering, we converted the images to black and white using thresholding. Subsequently, small distinct objects in the background of the mask images, mostly soil particles or fallen leaves of bright color, were removed by thresholding.

Step 4: We multiplied the mask image above and the raw (original) image, such that only the plant exhibited a color pixel value of more than 0, enabling the measurement of the average and median of RGB and L*a*b* values of the plant without background.

The images were processed in a Linux Ubuntu OS using eight CPUs with four cores (Intel Xeon Cache 10 MB, Intel (R) Xeon (R) CPU E5-1620 v4 @3.5 GHz) and 32 GB memory. Furthermore, we used a GPU (GeForce GTX 1080 Ti) with 11 GB graphic memory for computationally demanding tasks, including registration, augmentation, segmentation, and training using a DNN.

### Evaluation of the image analysis outcome

We prepared ground truth data (manually annotated ‘correct’ plant area) consisting of nine, nine, and twelve labeled single-plant images from the soil background at the Swiss site, sand background at the Swiss site, and sand background at the Japanese site, respectively. The images were selected, such that they represented variations in color, shape, and size among plants and backgrounds in all the images. Using the ground truth and outcome of the DANet, we calculated the Dice coefficient, precision, and sensitivity as follows:

- Dice coefficient = 2 × Area of overlap between ground truth and DANet outcome / total number of pixels in the ground truth and DANet outcome,
- precision = tp / (tp + fp),
- Sensitivity = recall = tp / (tp + fn),

where tp, fp, and fn indicate true positives, false positives, and false negatives, respectively.

### Anthocyanin and color of leaves

To examine how appropriately the leaf color represents the anthocyanin content, we photographed the leaves of plants growing in the common garden, harvested them, and quantified anthocyanin at the Swiss site, according to the absorption spectrometry method that detects wide molecular species of anthocyanin^71^. Relative anthocyanin content was calculated as (A_530_ nm – A_650_) / leaf area (mm^2^). We included all genotypes sampled on multiple dates to cover the wide anthocyanin accumulation levels. The detailed methodology is provided in **Supplementary Note 2**.

RGB images were acquired 150 cm above the plants using a RICOH WG-40 camera with the same setting as the time-series image acquisition (Table S6). Thereafter, we sampled target leaves to maintain them as frozen at −80 °C until anthocyanin extraction.

Before the analyses, the images were converted from the sRGB scale to the L*a*b* color space using the Python (ver 3.8.3) package grDevices.

Data analyses were conducted in R 4.1.0 (R Development Core Team 2021), unless otherwise stated. To examine the relationship between the relative anthocyanin content per leaf area and color information from images, we compared generalized linear models, linear models, and random forest models with anthocyanin content as the response variable and R, G, and B or L*, a*, and b* as explanatory variables^72^. The random forest model was developed using the *randomForest* package with default parameter settings. The details of the R packages used in this study are summarized in Table S7. We did not include the genotype or date effect in the model to avoid overfitting and to allow genotype- and date-nonspecific prediction of the pigment. We evaluated the accuracy of the models by leave-one-out cross-validation, where one out of the 451 data points was removed from the training dataset and used for prediction in each trial (Fig. S14) and by calculating the average of the root mean square errors for 451 trials (Table S4). We also conducted another validation in which a series of data points sampled from the same individual plant at different times were separated from the training dataset to avoid potential data leakage (Table S4). In both cases, the random forest model with L*a*b* performed the best. We plotted the validation result of this model and examined the relative contribution of a*, b*, and L* to the anthocyanin content using the ‘importance’ function in *randomForest*.

### Time-series plotting

To obtain a visual understanding of seasonal changes in plant traits, we plotted the plant area, L*, a*, b*, and estimated anthocyanin content in a time series. To enhance visibility, we plotted the 5-day moving averages of all plants of the same genotype per site per year using the packages *zoo*, *dplyr*, *ggplot2*, and *tidyverse*. We removed images from one camera at the Swiss site from yr2 because the plant IDs in the images could not be confirmed at the time of the analyses. In addition, we removed anomalous data according to [1] manual records of snow, storms, and other incidents that affect segmentation quality, [2] visual inspection of time-series plots for a*, where the cases of deviation from the norm were followed up based on the inspection of the original images to identify obstacles such as mesh and extreme cases of positioning failure, and [3] exclusion of the period of drastic value changes (outliers) in the time-series plots of the plant area. This was intended to filter out biologically unreasonable size fluctuations in the data and was performed via nearest neighbor imputation as follows: [3-1] Divide the time series into 10 blocks for each season for each site. [3-2] Mask one of the 10 blocks to be treated as a missing value. [3-3] Impute the missing values via the nearest neighbor method. [3-4] Calculate the difference between the imputed and original values. Referring to the score Z > 5 based on the difference, we identified and removed the data from 2018/02/18 – 2018/03/12 at the Swiss site that corresponded to the period of snow, frost, and strong light (Fig. S15). The analyses and plots were performed using Python ver. 3.9.10. The data_removed sheet in **supplementary Dataset 2** summarizes the dates on which all data were removed from the analysis as well as the reasons for removal. Generally, the initial plant area was large at the Swiss site and small at the Japanese site (Fig. S16). A small size increase during the season likely reflects the large starting size at the Swiss site, whereas at the Japanese site, it could be a combination of a real phenomenon with the difficulty in segmenting small plants (Fig. S16b).

### Dimension reduction with PCA

To compare the time-series trend of the estimated anthocyanin content, we compiled data on the genotype average of the estimated anthocyanin content from three seasons from two sites into one dataset. Following previous studies analyzing time-series data, we conducted dimension reduction by PCA using the package *base* and *vegan*^73–75^. To explore the time points that likely influenced the emerging pattern, we plotted the PC1 and PC2 scores over time using the R package *tidyverse*. To examine whether the trend was consistent between the two sites, we also conducted PCAs on the dataset in which the two sites were separated.

In addition, we investigated how well the average genotype represented the variation within the genotype. Accordingly, we split the dataset into sites and years, as the individuals were not consistent across sites or years. For each dataset, we performed Bayesian PCA using the *pcaMethods* package. This method complements missing values suitably when they constitute up to approximately 15% of the data, which is the case in our datasets (Fig. S12).

### Environmental data collection

To record environmental conditions, we set up weather stations at each site for precipitation, radiation, and air temperature. Precipitation data contained both rain and snow. The WS GP2 weather station (Delta-T Devices) was installed at the Swiss site, whereas at the Japanese site, different parameters were collected using devices from different suppliers (Table S8). The data were collected at 10 and 5 min intervals at the Japanese and Swiss sites, respectively.

Because of interruptions in data collection, precipitation and radiation data in the Swiss site were substituted by data from a nearby weather station of the Swiss Federal Office of Meteorology and Climatology (MeteoSwiss) in Fluntern. Missing values in air temperature data in the Swiss site were estimated using the *imputeTS* package.

### Relationship between estimated anthocyanin and environmental factors

To analyze how the estimated anthocyanin content is associated with environmental factors, we included air temperature, radiation, and precipitation as the key environmental factors in the analysis based on previous studies, field observations and laboratory experiments^22,45,53,76^ (Fig. S17). The threshold temperatures of 1, 4, 7, 10, and 13 ℃ were set for the calculation of coldness from air temperature to cover the range adopted or identified as critical in previous studies on cold treatment in *A. thaliana*^76–80^ (Fig. 6a). Considering the response time of plants to environmental cues, we determined for each environmental factor “reference windows,” i.e., durations for accumulating signals for anthocyanin content, and “lags,” i.e., durations between signal accumulation and anthocyanin content of a given day (Fig. 6a). The ranges of the reference windows and lags were 1–14 days and 0–14 days, respectively. We split the dataset into Swiss and Japanese sites to account for the differences in climate and the trend of fluctuation in the estimated anthocyanin content. For each genotype, we fitted linear regression models with coldness, radiation, and precipitation as explanatory variables after standardization and estimated anthocyanin content as a response variable using one of the 46,305,000 parameter combinations (5 temperature thresholds × 14^3^ windows × 15^3^ lags). We selected the model with the smallest Akaike information criteria as the best model and used it to examine the association between environmental factors and estimated anthocyanin content. These calculations were performed using R. 4.2.0. As the assumptions of the linear model were not always met, we determined the significance of the explanatory variables by calculating confidence intervals with the bias-corrected and accelerated bootstrap in the *boot* package. The relative contributions of the explanatory variables were extracted using the *relaimpo* package. In addition, we generated time-series plots of the estimated anthocyanin content and environmental factors with the best parameters for the model plant *A. thaliana* using the *ggplot2* and *reshape2* packages.

## Data availability

The images and labelling data obtained and analyzed in the study are deposited in Dryad. The remaining data are available from the corresponding authors upon request.

## Code availability

The scripts used for the analyses of the images and data collected from the field and the data qgenerated therefrom can be freely and openly accessed at Zenodo. The remaining scripts are available from the corresponding authors upon request.

## Acknowledgements

The authors thank Dennis Bailer, Richard Baxter, Gioele Bello, Silas Braun, Silvana Capaul, Mizuki Sato, Marcel Freund, Matthias Furler, Anne Graf, Nangsa Karutshang, Lucas Mohn, Aki Morishima, Atsushi J. Nagano, Kenji Ogawa, Elin Rütimann, Yasuhiro Sato, Reinhold Stockenhuber, Karen Thomsen, Misako Yamazaki, Mayumi Yorino, Center for Ecological Research of Kyoto University, and Kudoh Lab at Kyoto University. This study was funded by Core Research for Evolutionary Science and Technology grant number JPMJCR16O3 to KKS, YS, JSese and TK, JPMJCR15O1 to HK, Swiss National Science Foundation grant 31003A_182318, CRSII5_183578, Kakenhi 22H02316 to KKS and TT, JP21H04977 to HK, University Research Priority Program in Evolution in Action to KKS and RSI, 22H05179 to KKS and JSun, and Joint Usage/ Research Grant of Center for Ecological Research (2022jurc-cer02), Kyoto University to RSI.

## Authors’ contributions

RA, RSI designed the experiment, RA, TT, JSugisaka, and RSI conducted the experiment, RA, TG, TT, MH, KK, and JSun analyzed the data, RA, TG, TT and KKS wrote the manuscript, JA, NK, and JSese provided technical support, and YS, JSese, NK, KKS conceived the study. All authors contributed to revising the manuscript and approved the final version.

---

**Supplementary Movie 1**: Time-series compilation of the segmented plant area

Images of *Arabidopsis thaliana* (COL) from the Swiss site during the 2018–2019 season (yr2). The cyan lines indicate the segmented areas. The corresponding RGB values are shown in the images. Anomalous data were removed as described in the Methods section, except for snow cover that was retained to facilitate the visual recognition of the effect of snow on the target plants.

ZHA04-COL-51ContDatetimePlotRGBMJPGxdate_subframes1.avi

**Supplementary Movie 2**: Time-series compilation of the segmented plant area

Images of Japanese *Arabidopsis kamchatica* (MAG) from the Swiss site during the 2018–2019 season (yr2). The cyan lines indicate the segmented areas. The corresponding RGB values are shown in the images. Anomalous data were removed as described in the Methods section, except for snow cover that was retained to facilitate the visual recognition of the effect of snow on the target plants.

ZHA04-MAG-76ContDatetimePlotRGBMJPGxdate_subframes1.avi

**Supplementary Movie 3**: Time-series compilation of the segmented plant area

Images of synthetic *Arabidopsis kamchatica* (RS7) from the Swiss site during the 2018–2019 season (yr2). The cyan lines indicate the segmented areas. The corresponding RGB values are shown in the images. Anomalous data were removed as described in the Methods section, except for snow cover that was retained to facilitate the visual recognition of the effect of snow on the target plants.

ZHA04-RS7-77ContDatetimePlotRGBMJPGxdate_subframes1.avi

**Supplementary Note 1:** Plant Cultivation

Swiss site

Seeds of *Arabidopsis lyrata*, natural and synthetic *A. kamchatica*, and *A. thaliana* were sown in 12 well plates (Nunclon™ Delta Surface, Thermo Scientific, Denmark) with quartz sand (0.1–0.8 mm TOP MINERAL AG, Switzerland) and placed at 4 °C for a week and subsequently by a window to stimulate germination. The seedlings at the cotyledon stage were transferred to biodegradable pots (W × L × H: 3 cm × 3 cm × 5 cm Peat Pot Strips, Jiffy) filled with a mixture of Floratorf (Floragard, Oldenburg, Germany): quartz sand (0.4–0.8 mm Quarzsand, Carlo Bernasconi AG, Switzerland) = 1:1 in volume. We blended ActaraG (Syngenta Agro, Switzerland) with this soil mixture. The potted seedlings of *A. lyrata* and *A. kamchatica* were cultivated in a growth chamber with a long-day setting (22 °C/20 ℃, 16 h:8 h light: dark, RH 60%, light 120–140 μE) for six weeks, while *A. thaliana* seedlings were placed in a short day chamber for three weeks to keep them vegetative (18 °C/16 ℃, 8 h:16 h light: dark, RH 60%, light 120–140 μE). For *A. halleri*, we cultivated clonally propagated branch segments in pots in a long day chamber for five weeks. For all species, potted plants were placed on a plastic tray covered with a transparent lid that was half-opened on the 3rd day and fully opened one week after potting. At potting and once per week, watering was performed using Wuxal Universaldünger nutrient solution (Maag, Westland Schweiz GmbH, Switzerland). We acclimated all plants outside under the roof for a week before transplanting them to the common garden at the Irchel Campus of the University of Zurich. The seedlings were planted in two built-in compartments, each 1 × 7 m, filled with well-watered Rasenerde (Ökohum GmbH, Switzerland). Only in yr3 did we have a ca. 5 cm layer of quartz sand on the soil surface to ensure better contrast between plants and the background. The ground was watered immediately before transplanting the seedlings. The entire compartment was covered with polyester mesh sheets (1 mm × 1 mm grid) over the skeleton of the polytunnel (model A-17, Nan-ei Kogyo, Japan). During winter, the mesh at the top was removed to allow snow fall and prevent the skeletons from collapsing owing to strong winds.

The Japanese site

The seeds of *A. lyrata*, natural and synthetic *A. kamchatica*, and *A. thaliana* were sown on sterilized plastic dishes (Asnol Petri Dish φ90 × 20mm, 1-8549-04, AS ONE Corporation) filled with ca. 60 g of 0.3-0.6 mm quartz (#5, Toyo Matelan Co. Ltd.) dampened with tap water. The dishes were kept in an incubator (KOITOTRON HNM-S11, KI Holdings Group) at 21℃/15℃, 12h:12h light: dark for four weeks, except for *A. thaliana* which was kept there for one week. When the seeds did not germinate after four weeks, the dishes were kept in the chamber for up to additional two weeks. The seedlings were planted on blocks of mineral wool (rock fiber for cultivation M40T40, Nittobo) that were kept in an incubator (22°C/20℃, 16h:8h light: dark, RH 74%, KOITOTRON HNM-S11) for six weeks, except the branch segments of *A. halleri* prepared in the same manner as in the Swiss site and the seedlings of *A. thaliana* which were brought in one week later and three weeks later, respectively. Watering was performed twice a week, once of which using × 2000 HYPONeX solution (HYPONeX Japan Corp., Ltd.). All seedlings were acclimated for a week outside under the roof. The seedlings were transplanted in a compartment (W × L × H: 100 cm × 100 cm × 20 cm) filled with a mixture of 40 L of humus (100% Shinshu Fall Leaves 100% Natural Fermented Products, Koshin Kawara Co. Ltd.) and 60 kg of the 0.3-0.6 mm quartz. We covered the surface of the soil mixture with a 1-2 cm thick layer of 0.3-0.6 mm quartz. The ground was wet immediately before transplanting and watered once directly after transplanting to promote the seedling establishment. During the growing season, we set up a cage with mesh (20µm x 20µm, Moritaya, Japan & Marushin, Japan) over each compartment to prevent cross-pollination among experimental plants. A root-cutting-sheet (‘Kurapapy’, Kuraray Co. Ltd. in yr1 and yr2 and ‘Paopao Nekiri Sheet’ Nihon Nougyou System in yr3) was placed at the bottom of the soil to prevent overgrowth.

**Supplementary Note 2:** Anthocyanin content measurement and calculation of relative anthocyanin level

We photographed the targeted leaves of plants growing in the common garden, harvested them, and quantified anthocyanin content by the method of absorption spectrometry^1^. These plants were grown in addition to the plants for image acquisition. In yr2, one leaf per plant was subject to the study for DEN, RS7, and RS8. We had eight replicates per genotype at each of the four time points (October 31, November 20, February 14, and March 20). At each time point, we used plants that were not subject to the study at the earlier time point(s). In yr3, one leaf per plant from the remaining nine genotypes (Cf.) Table S1) was subject to the study at each of the five time points (October 31, November 14, December 3, February 25, and April 23). We had eight replicates per genotype per time point. Due to limitations in space and the number of seeds, we used the same plants on October 31, December 3, and February 25, and on November 14 and April 23. The collected leaves were kept frozen at −80°C until processed. Each leaf was ground in 1000 uL of extraction buffer (18% isopropanol and 1% HCl) and incubated at room temperature in a shaded condition for 24 hours. After the centrifugation of 10 min at 15000 *g,* the absorbance of the supernatant was measured at 535 nm and 650 nm. The relative anthocyanin content was calculated by dividing the value (A_535_-A_650_) by the leaf area (cm^2^). The leaf area was measured from the images using Fiji ver. 1.52n^2^.

**Fig. S1.**
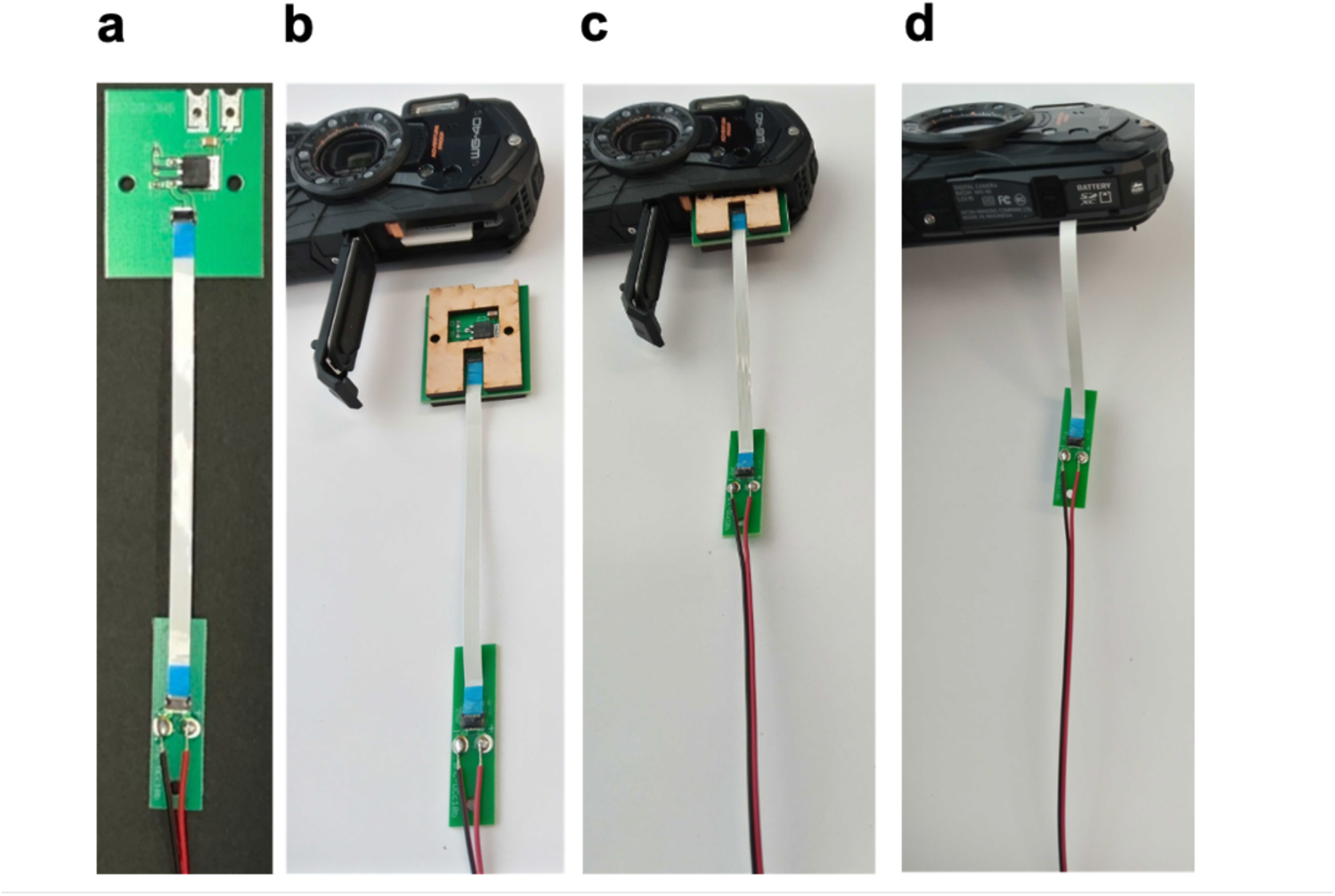
Details of the set-up consisting of a DC coupler, wooden fixing panels, a flat cable, and a connector. **a** The set-up without wooden fixing panels where the DC coupler is visible. **b** The set-up and a camera. **c** The DC coupler and wooden fixing panels being inserted in the battery slot of a camera. **d** The DC coupler and wooden fixing panels in the battery slot, while the flat cable coming out with the camera lid closed. The blueprint of the DC coupler is available at: https://github.com/akita11/RicohCameraDCCoupler

**Fig. S2.**
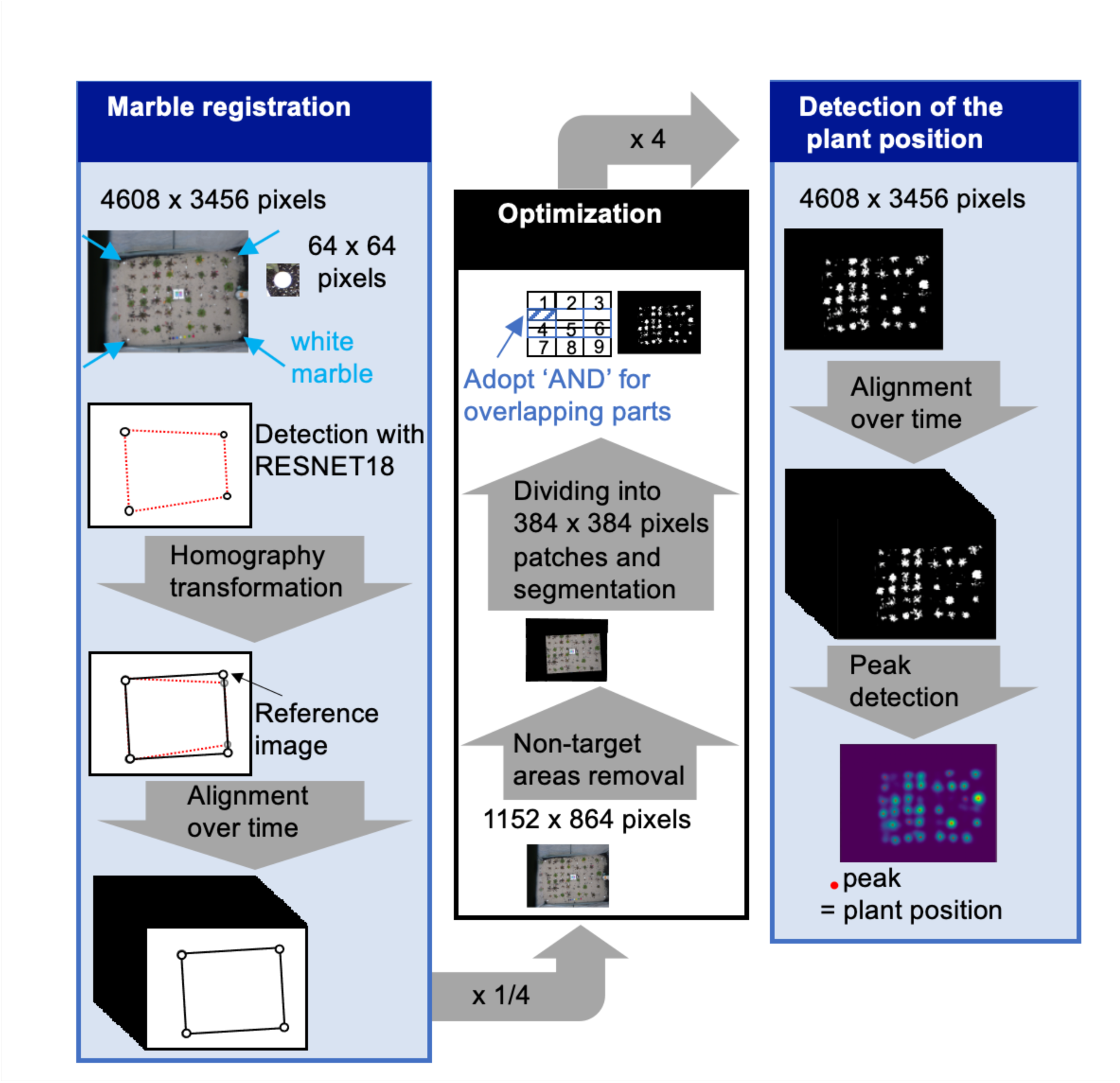
The workflow of the detection of the plant position. Marble registration was conducted first to align the position of the images over time. Optimisation was performed to reduce the processing time. The plant position was defined as the peak detected in the aligned images.

**Fig. S3.**
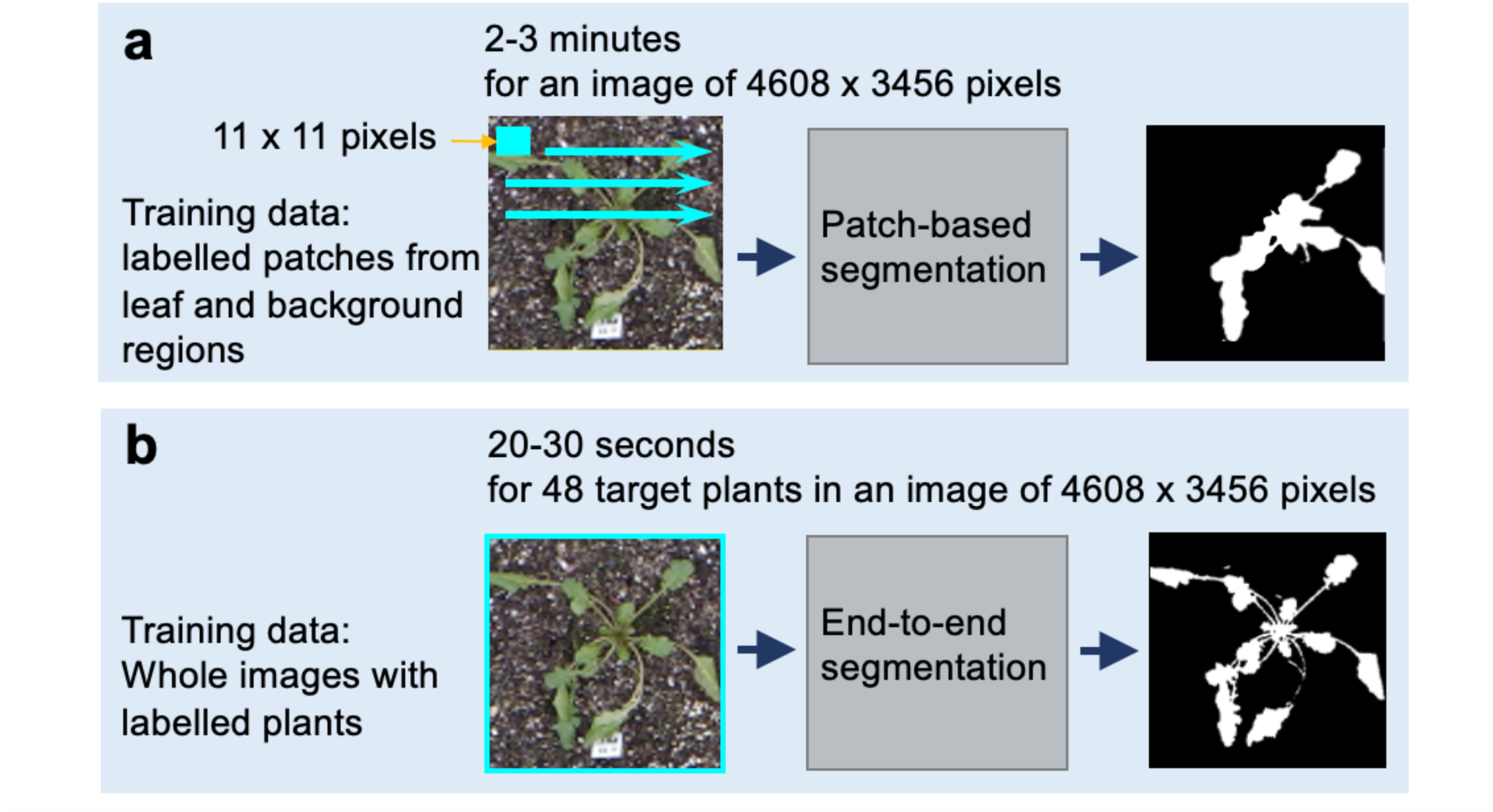
Workflows of **a** A patch-based and **b** an end-to-end segmentation. **a** Patchbased method: whether the pixel in the center of a patch of 11 x 11 pixels is a leaf or not was determined. This procedure was repeated for all the 4608 x 3456 pixels in the image. **b** End-to-end method (DANet): an area including a target plant was used as an input.

**Fig. S4.**
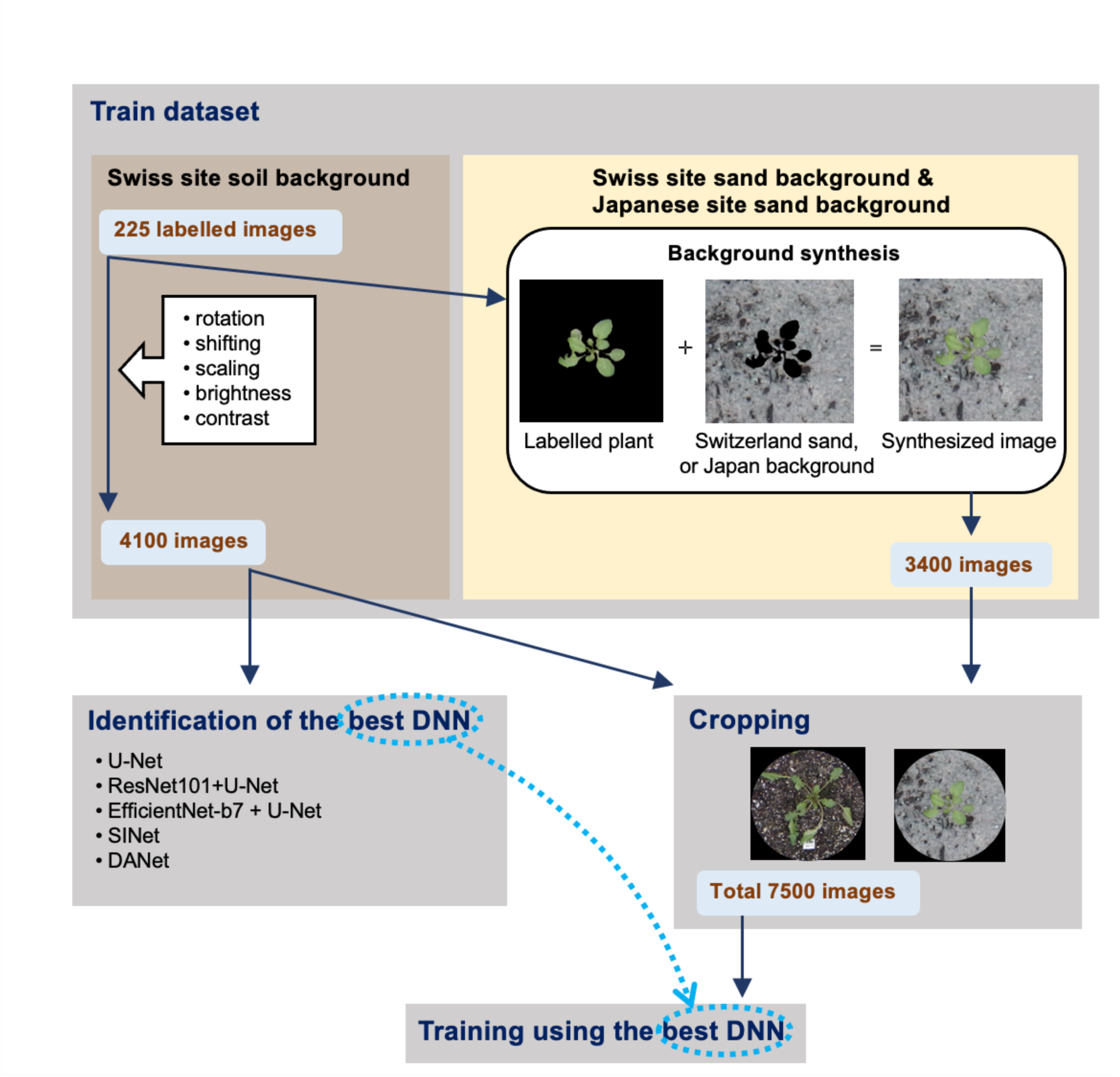
Training of the plant segmentation model, consisting of the preparation of the train dataset, identification of the best **DNN** for the study dataset, and cropping of the images for training. Processes with white background correspond to augmentation.

**Fig. S5.**
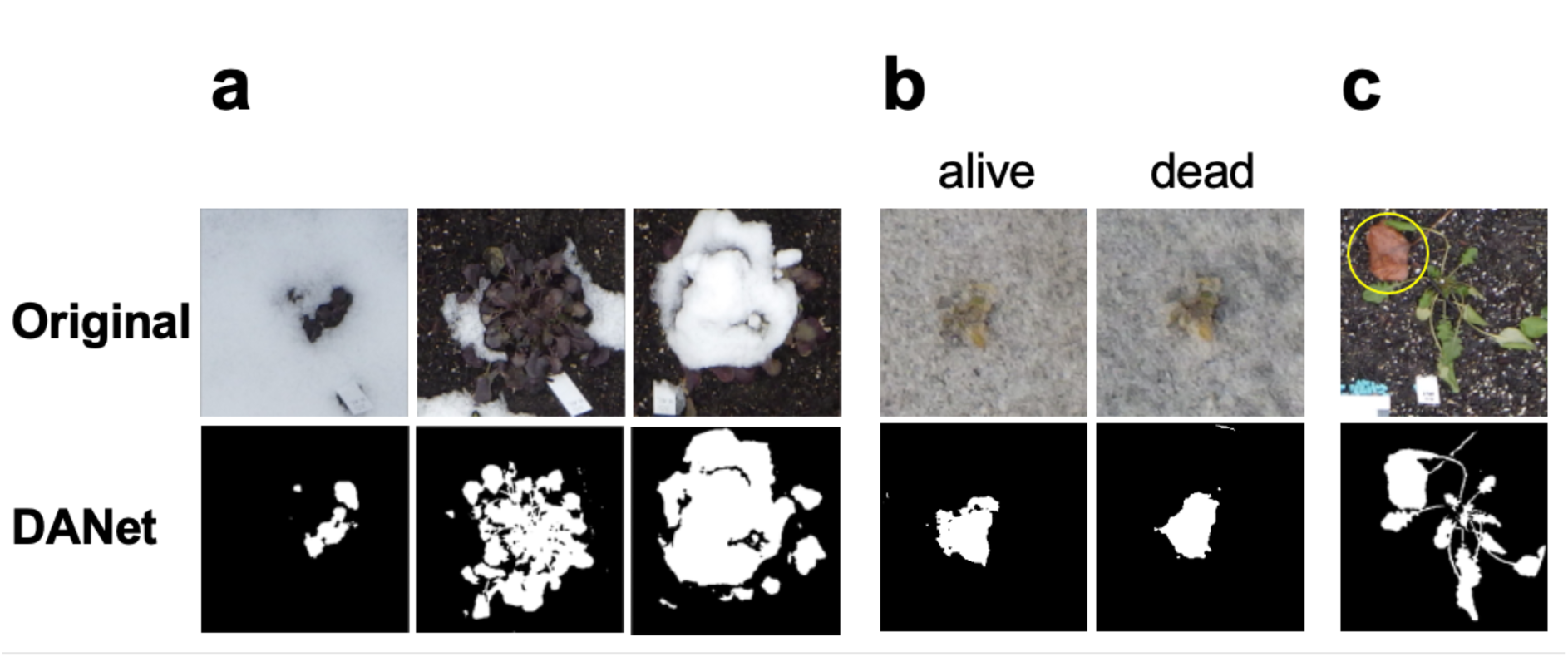
Examples of factors complicating the interpretation of the segmentation outcome. **a** snow. **b** plant death. **c** fallen leaf (in yellow circle). In **a-c**, the original image and the outcome of the segmentation by DANet are shown in the top and bottom rows, respectively.

**Fig. S6.**
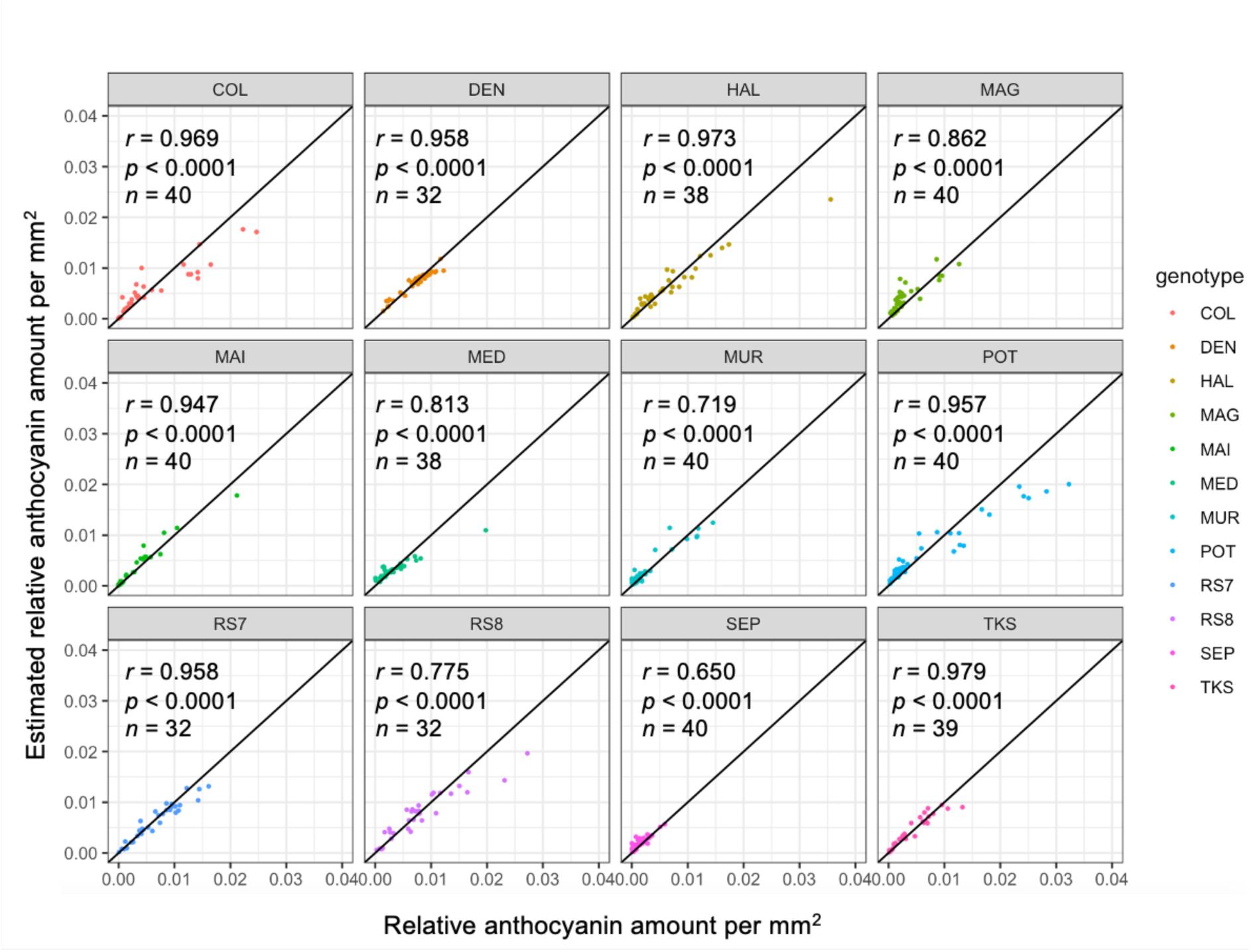
The results of fitting of the random forest model to estimate the relative anthocyanin per mm^2^ for each genotype. The scatter plots show the correlation between the relative anthocyanin per mm^2^ and the estimated relative anthocyanin per mm^2^ using the random forest model. Correlation coefficients (*r*) and p-values are indicated in the plots.

**Fig. S7.**
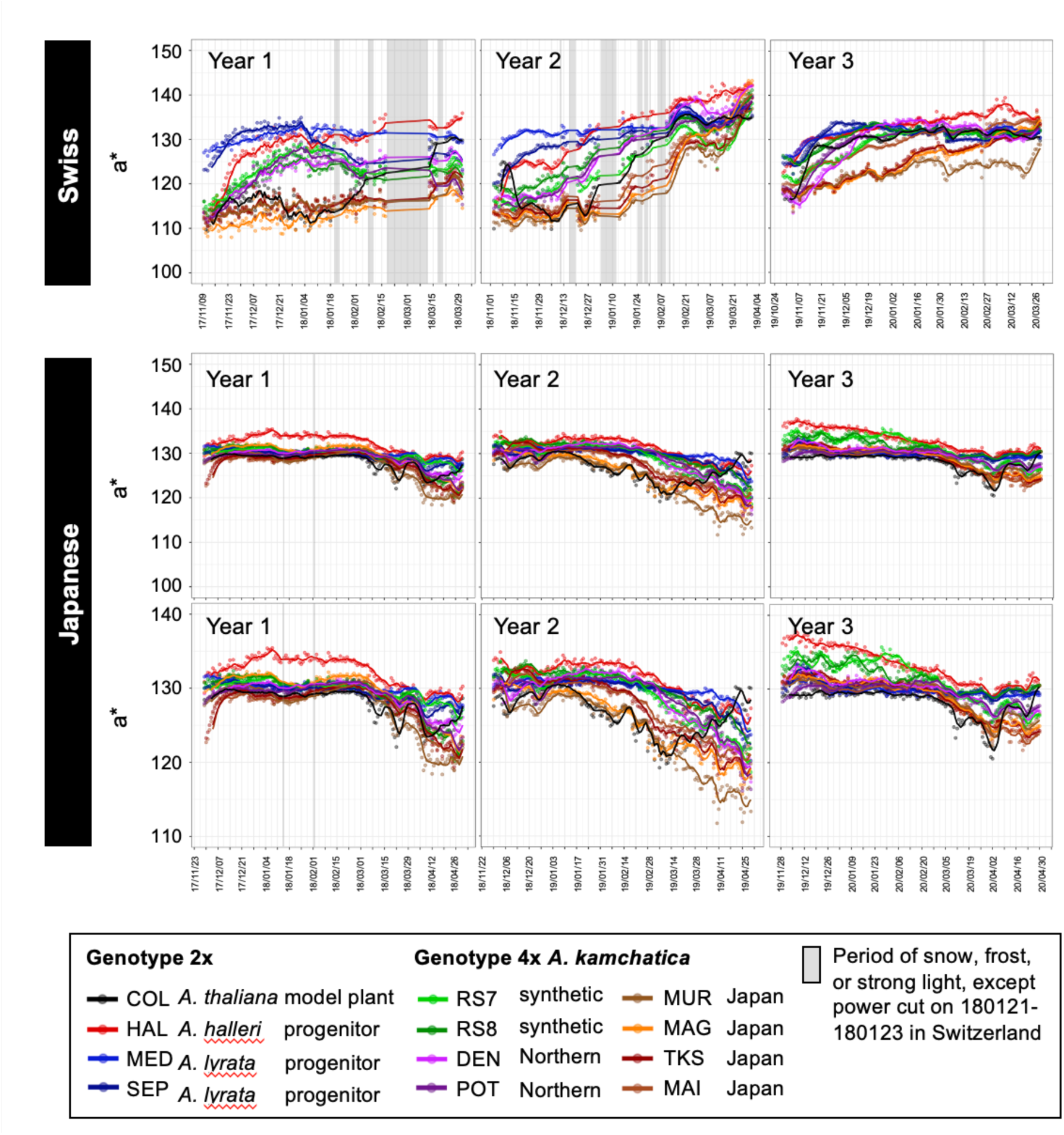
Time-series plots of 5-day moving average of a* in plants of 12 *Arabidopsis* genotypes at the Swiss and Japanese sites in three seasons. The plots of the Japanese site are shown in two versions, where the top row presents the same Y-axis scale as that of the plots of the Swiss site and the bottom row presents the plots with enlarged Y-axis for better visibility.

**Fig. S8.**
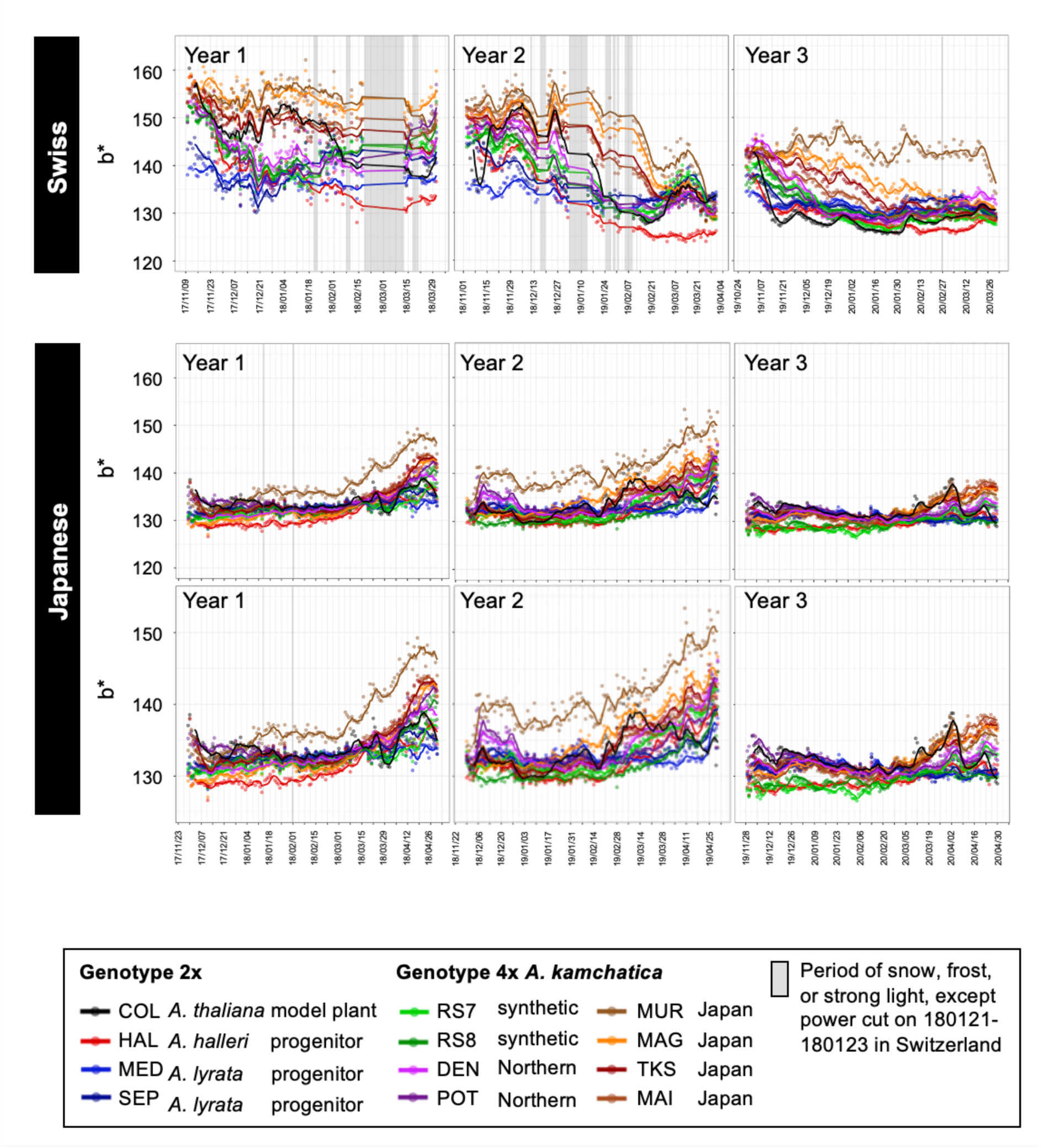
Time-series plots of 5-day moving average of b* in plants of 12 *Arabidopsis* genotypes at the Swiss and Japanese sites in three seasons. The plots of the Japanese site are shown in two versions, where the top row presents the same Y-axis scale as that of the plots of the Swiss site and the bottom row presents the plots with enlarged Y-axis for better visibility.

**Fig. S9.**
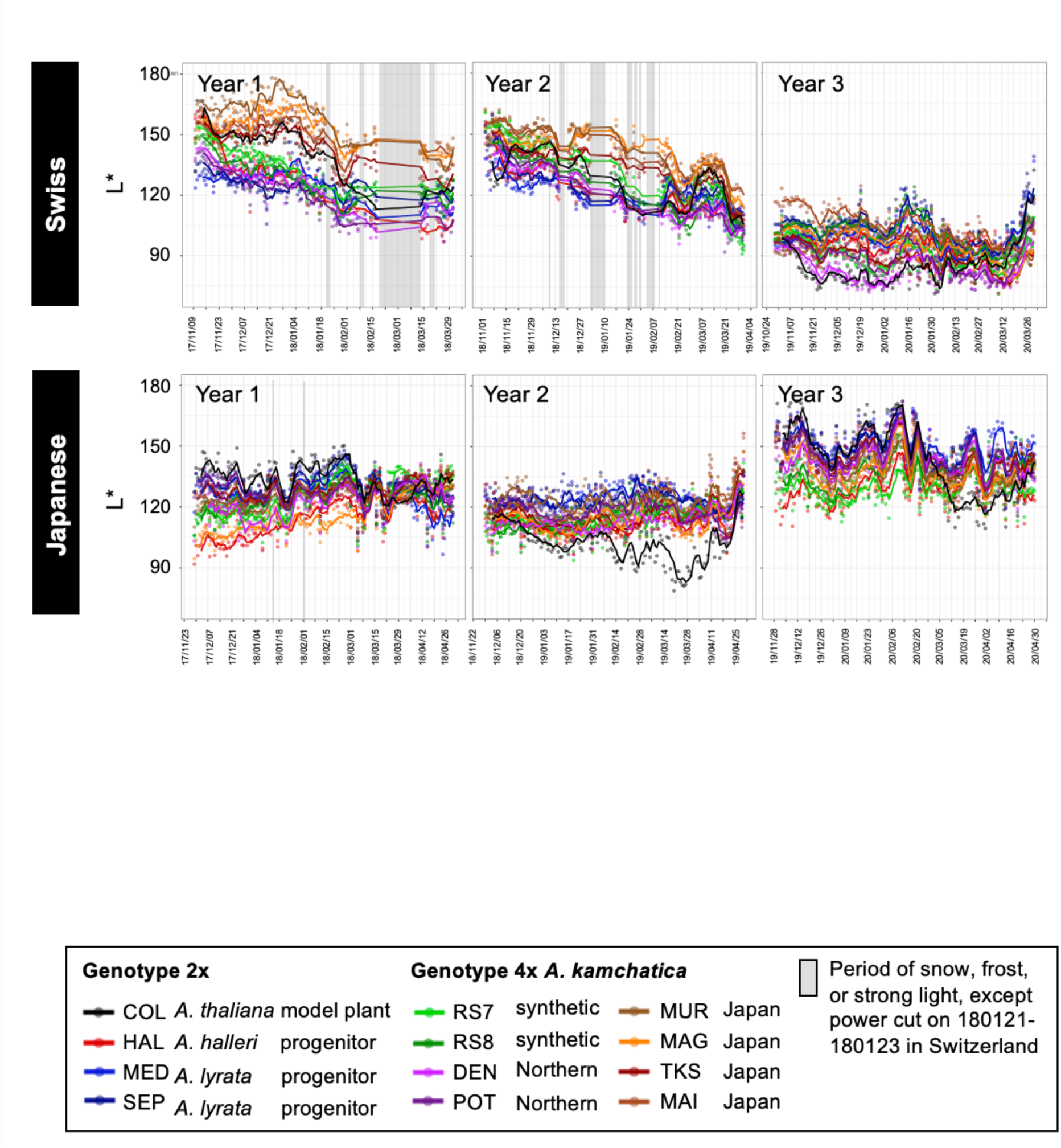
Time-series plots of 5-day moving average of L * in plants of 12 *Arabidopsis* genotypes at the Swiss and Japanese sites in three seasons.

**Fig. S10.**
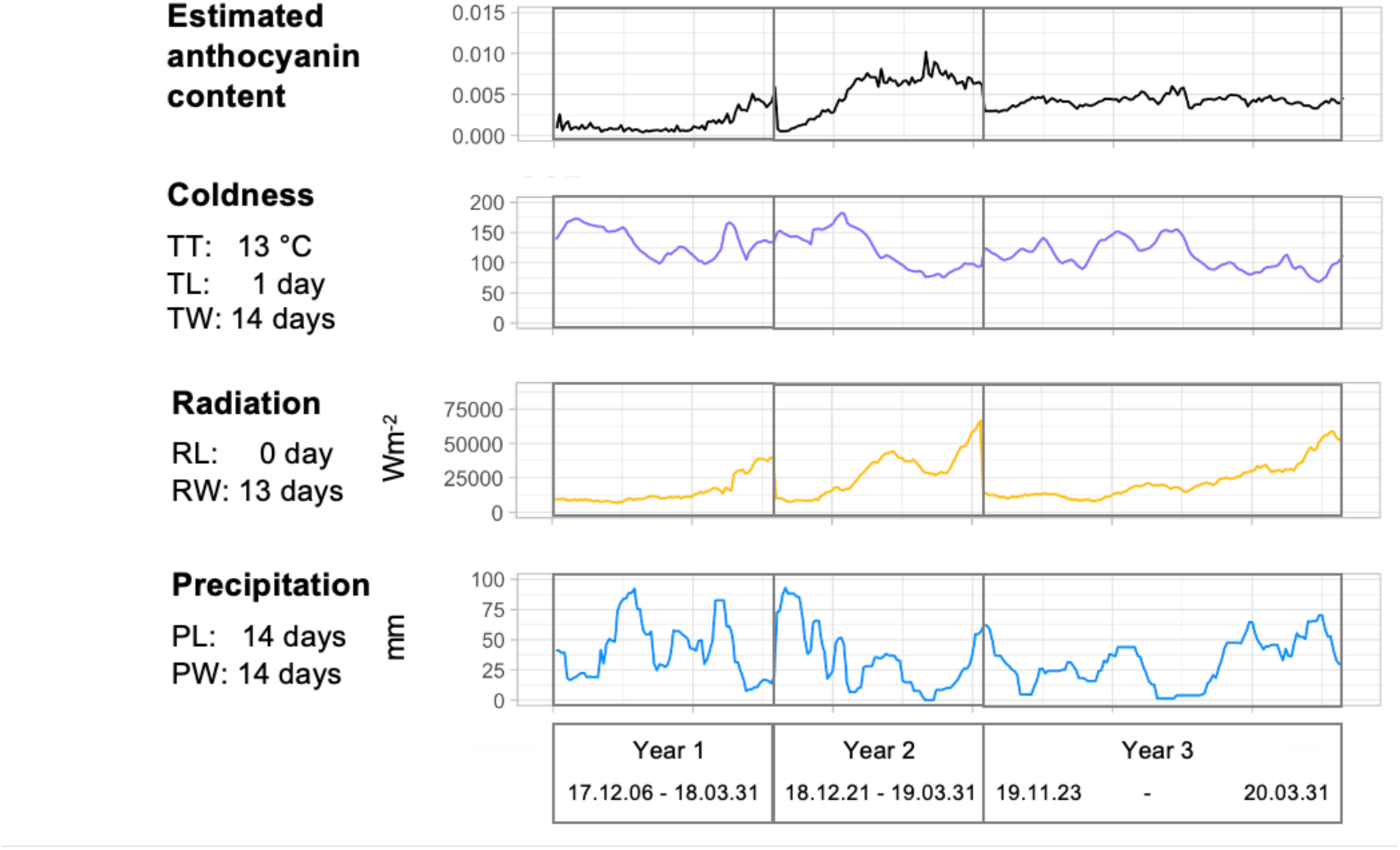
Time-series plots of the estimated anthocyanin content and the environmental variables with best parameter combinations for the model plant *Arabidopsis thaliana* in three years in the Swiss site. TT: temperature threshold, TL: temperature lag, TW: temperature window, RL: radiation lag, RW: radiation window, PL: precipitation lag, PW: precipitation window. See Fig.7a for the calculation of the environmental variables using the parameters. *n* = 663.

**Fig. S11.**
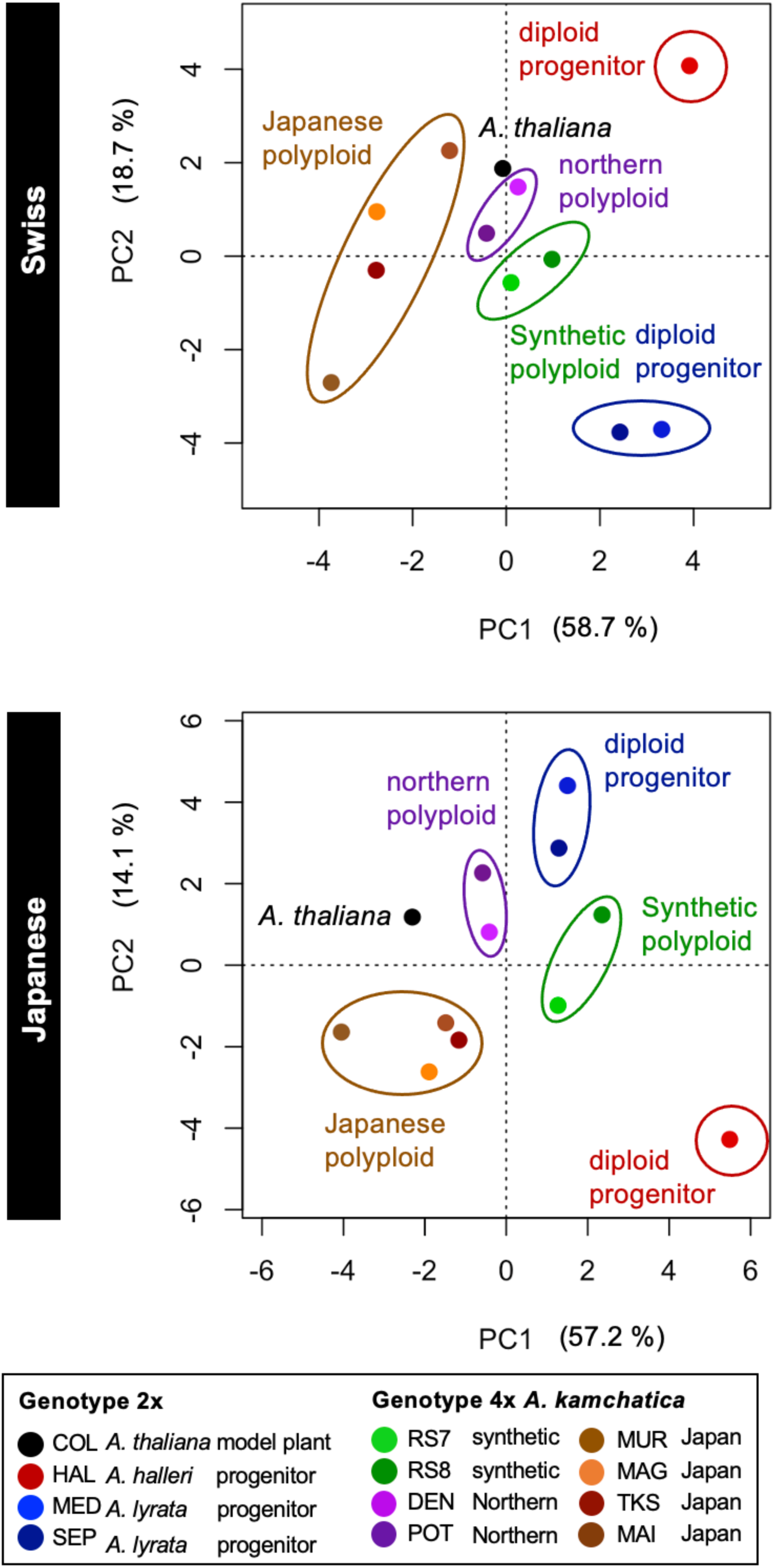
Principal component analysis plots of average estimated relative anthocyanin content per mm^2^ for 12 *Arabidopsis* genotypes. The data from three seasons are merged for Swiss and Japanese sites, respectively.

**Fig. S12.**
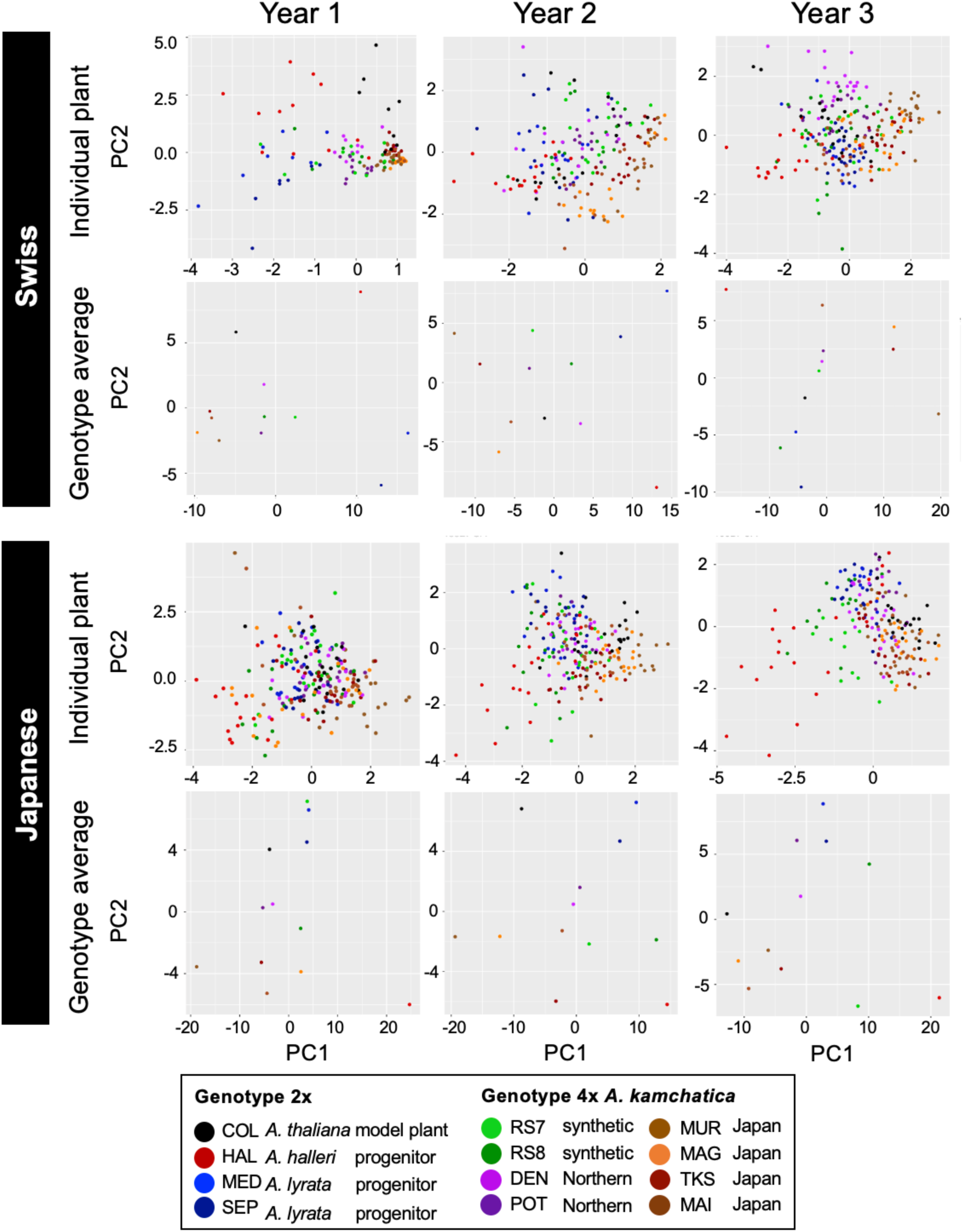
PCA plots for estimated anthocyanin content at Individual plant level and of genotype average for Years 1, 2, and 3 at the Swiss and Japanese sites. Because new sets of plants were grown each year and therefore the individuals were not consistent across years or country, year and country are separately plotted. Missing values at individual level data were imputed using Baysian PCA. The proportion of missing values was 10.1%, 11.7%, and 12.9% for Switzerland Years 1, 2, and 3, and 4.4%, 6.0%, and 4.0% for Japan Years 1, 2, and 3, respectively.

**Fig. S13.**
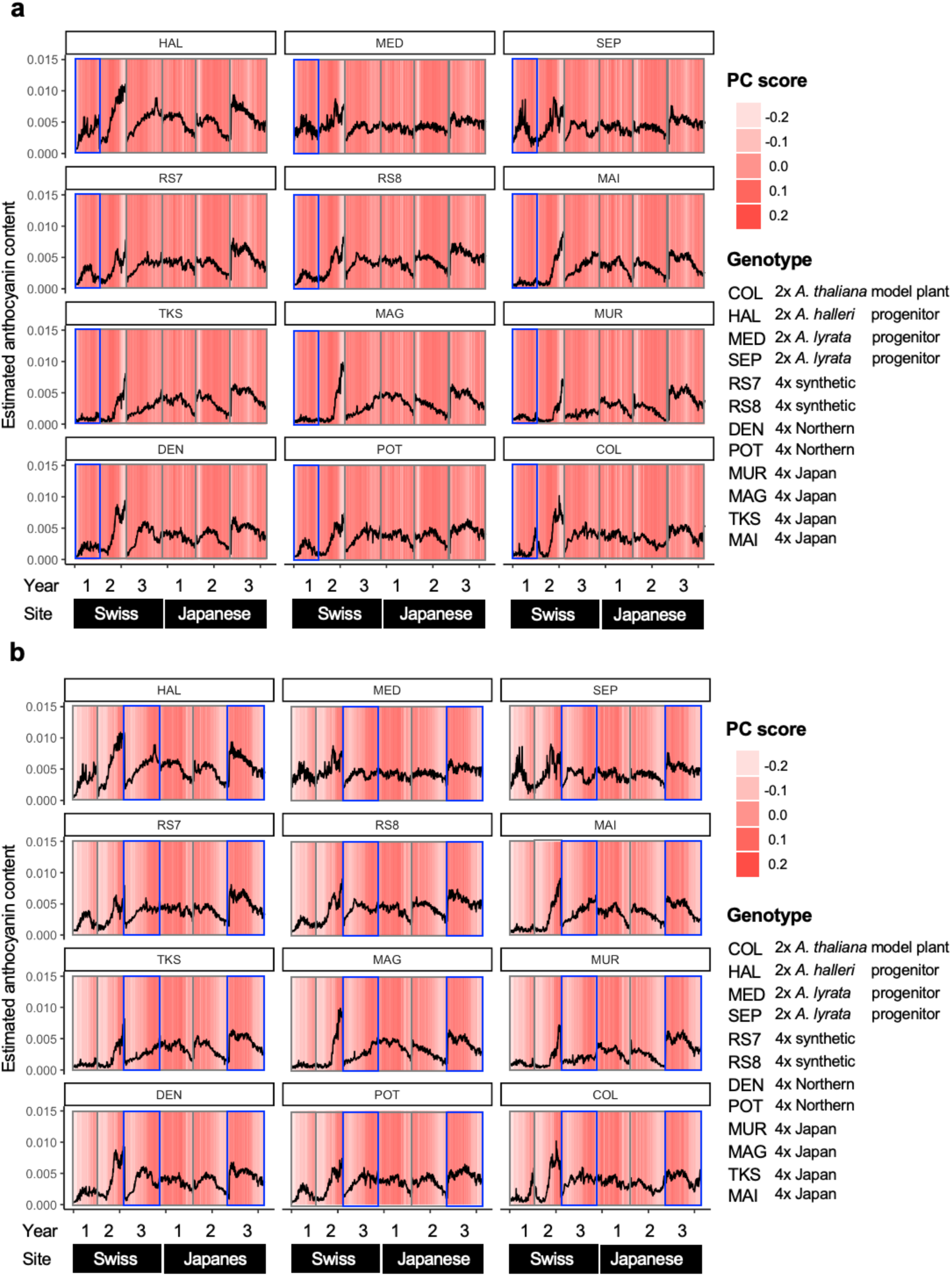
Time-series plots of the estimated anthocyanin content for 12 genotypes of *Arabidopsis* overlaid on a PC1 scores and b PC2 scores. 2x and 4x indicate diploid and allotetraploid, respectively. Blue rectangles indicate the period with distinct difference among genotypes and high PC scores. In addition, PC2 score seemingly correlated with the estimate anthocyanin content throughout the years and sites.

**Fig. S14.**
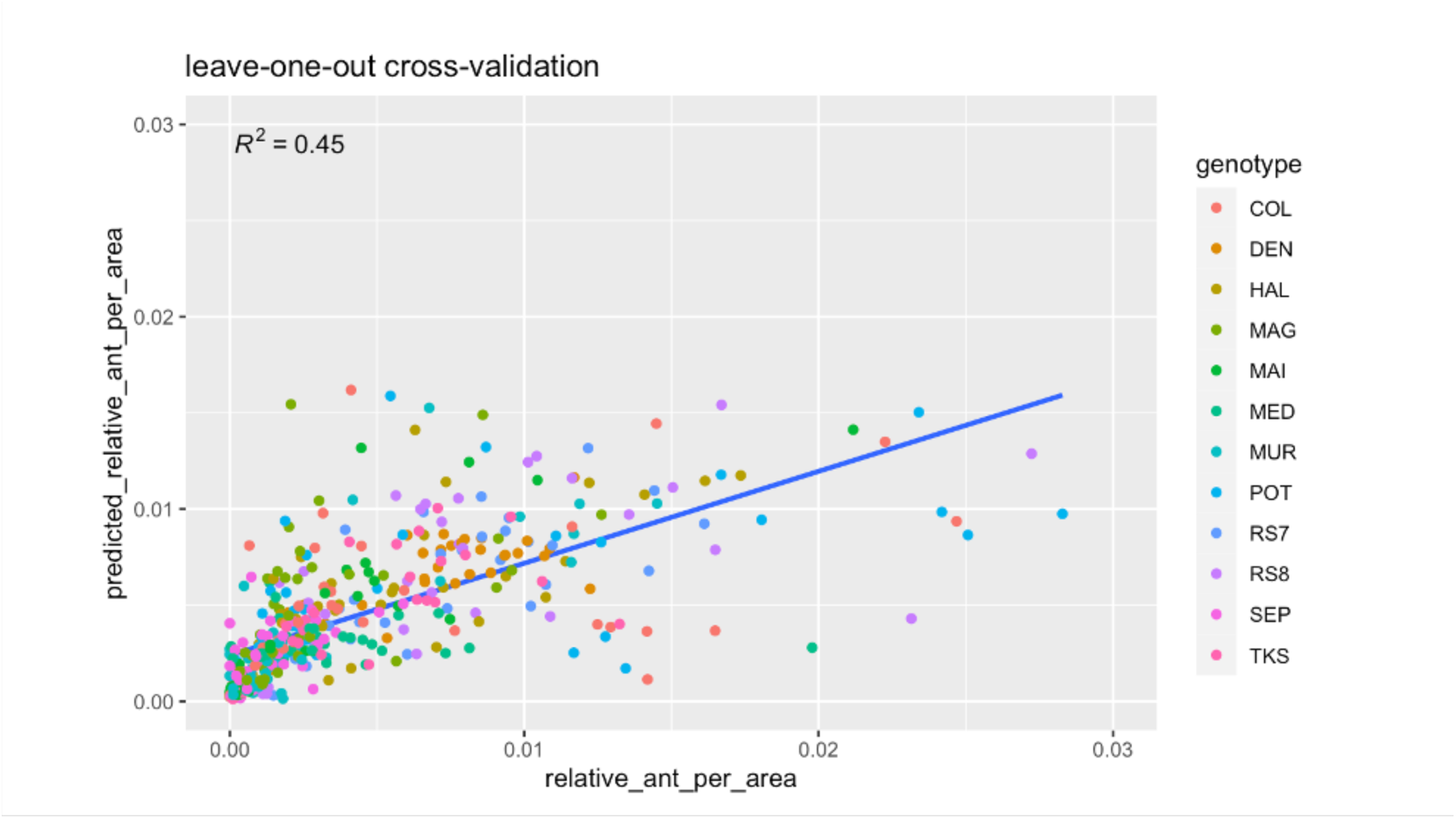
The result of the leave-one-out cross-validation of the random forest model to estimate the relative anthocyanin per mm2. The scatter plot shows the relation between the relative anthocyanin per mm^2^ and the estimated relative anthocyanin per mm^2^ when one data point is excluded per time using the random forest model. R-squared (R^2^) is indicated in the figure. *n* = 451.

**Fig. S15.**
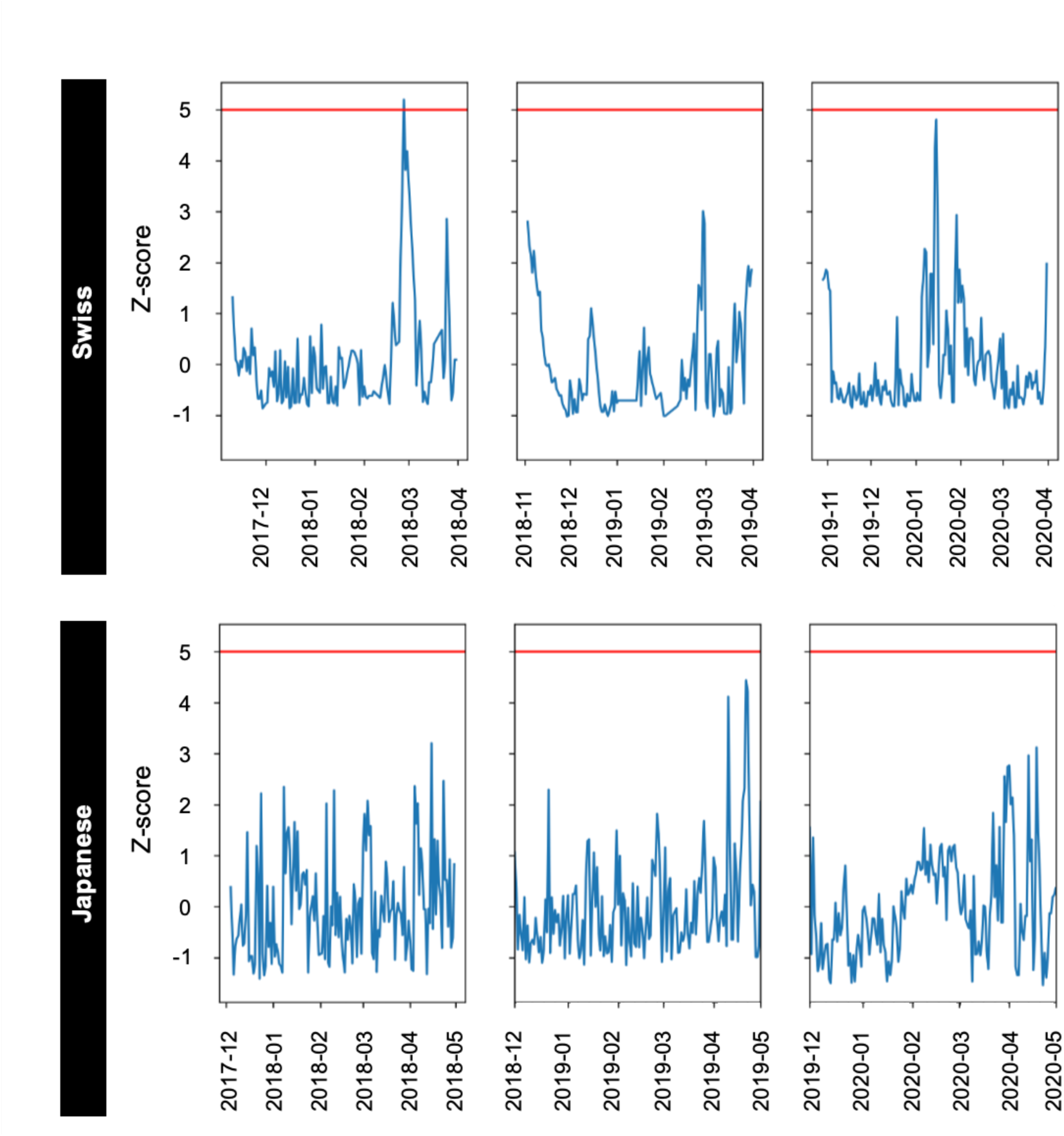
Detection of the period of drastic value changes (outliers) in the in time-series plots for plant area at the Swiss and Japanese sites in three seasons using nearest neighbor imputation.

**Fig. S16.**
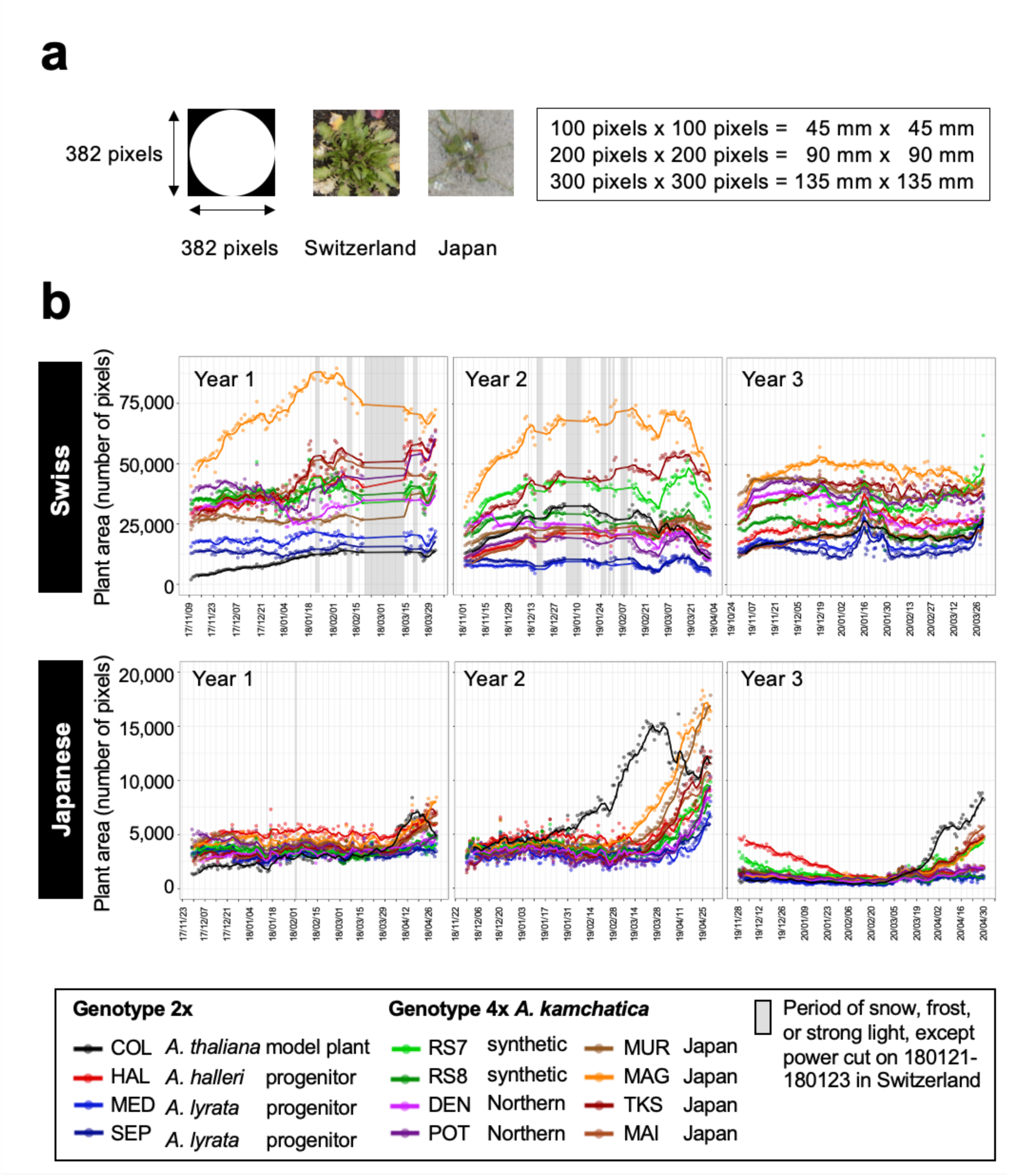
The plant area of *Arabidopsis* genotypes at the Swiss and Japanese sites. **a** The details of the measurement in pixels with representative images and the conversion to the area in mm^2^. **b** Time-series plots of 5-day moving average of the plant area of 12 genotypes at the Swiss and Japanese sites in three seasons. Note theY-axis scale difference between sites.

**Fig. S17.**
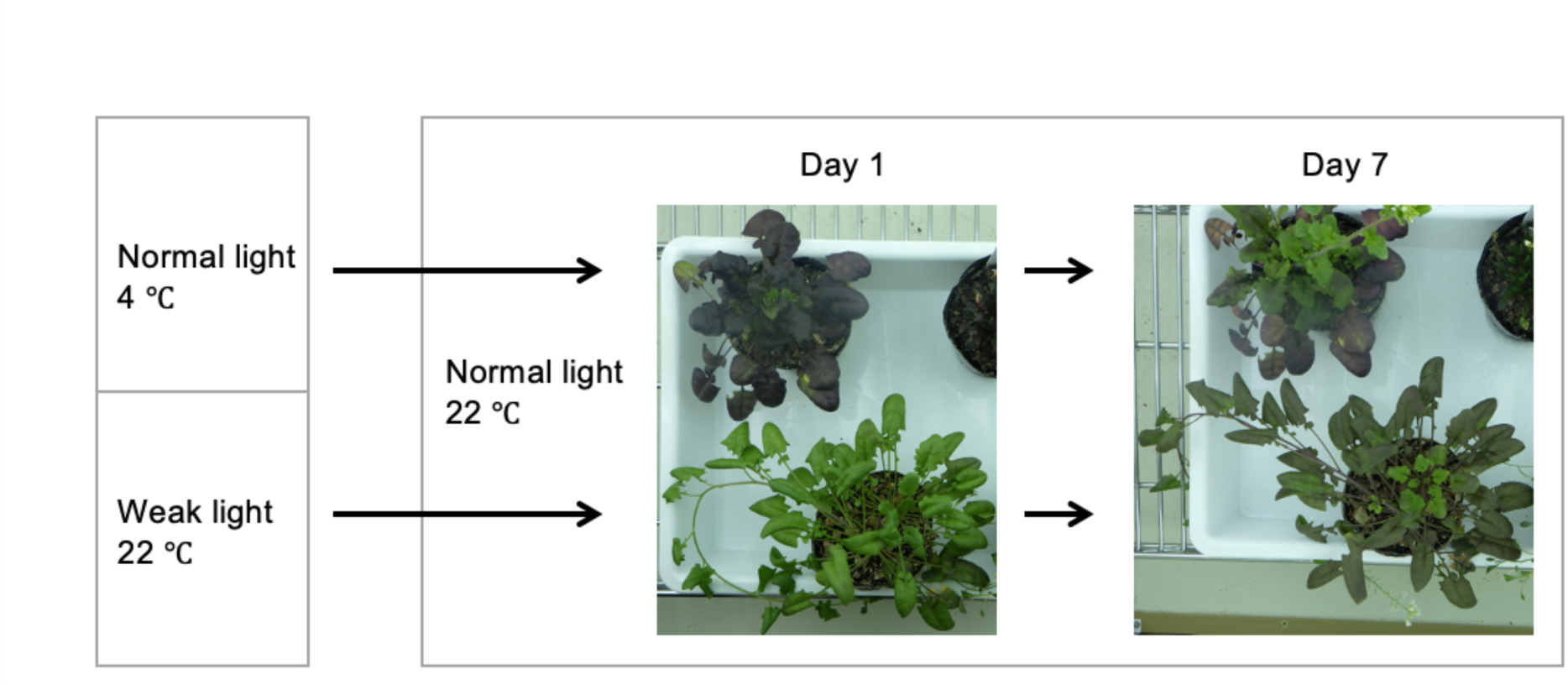
Images of individuals of *Arabidopsis halleri* W302 subject to experiments in which temperature and light intensity were manipulated. The plant above was first kept at low temperature and then brought to benign temperature under normal light condition (ca. 100 μmol m^−2^ s^−1^) to examine the effect of temperature on plant color. The plant below was first kept at weak light (ca. 40 μmol m^−2^ s^−1^) and then brought to normal light under benign temperature to examine the effect of light intensity on plant color. The day length was kept at short day condition throughout the experiment with 10 h: 14h = light:dark.

**Table S1.**
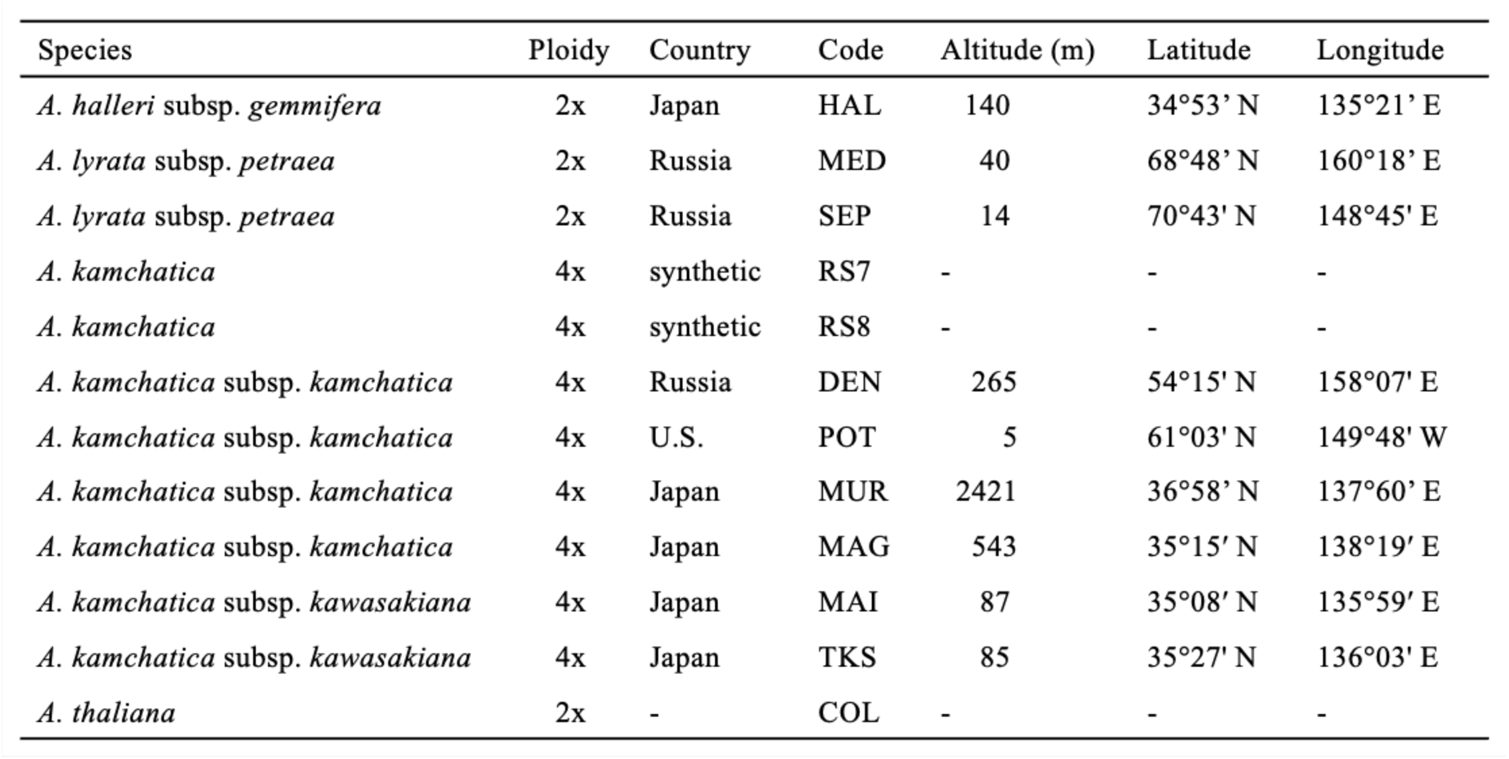
The name of the species, ploidy, genotype code, altitude of origin, latitude of origin, and longitude of origin of the genotypes of *Arabidopsis* used in the study. RS7 and RS8 were synthesized in the laboratory and thus have no information on the altitude, latitude, or longitude of origin. We used the lab accession Col-O for *A.thaliana*.

**Table S2.** The summary ofDice of the outcome of segmentation using different DNNs on 4,100 images of soil background from the Swiss site.

| Category | U-Net | Resnet101 + U-Net | Effectivenet-B7 + U-Net | SINet | DANet |
| --- | --- | --- | --- | --- | --- |
| average | 0.662 | 0.778 | 0.808 | 0.673 | 0.819 |
| minimum | 0.186 | 0.513 | 0.544 | 0.294 | 0.562 |
| maximum | 0.919 | 0.952 | 0.952 | 0.889 | 0.962 |

**Table S3.** The summary of assessment indices of the outcome of segmentation using DANet on 7,500 images from the Swiss (soil and sand backgrounds) and Japanese sites.

| Category | Dice | Sensitivity | Specificity |
| --- | --- | --- | --- |
| mean | 0.5767 | 0.8655 | 0.9544 |
| standard deviation | 0.2441 | 0.1292 | 0.0190 |
| maximum | 0.9098 | 0.9912 | 0.9804 |
| minimum | 0.1077 | 0.4760 | 0.9150 |

**Table S4.** Root mean square error of different types of models to estimate the anthocyanin content from color information (RGB, or L*a*b* color space) using two methods. Method 1 corresponds to Leave-One-Out Cross-Validation. Method 2 is a cross validation where data from the same individual plants were separated from training data set.

| Model type | Explanatory variables | Method 1 | Method 2 |
| --- | --- | --- | --- |
| linear | R + G + B | 0.00447 | 0.00445 |
|  | L* + a* + b* | 0.00442 | 0.00440 |
| generalised linear | R + G + B | 0.00464 | 0.00476 |
|  | L* + a* + b* | 0.00507 | 0.00507 |
| random forest | R + G + B | 0.00427 | 0.00425 |
|  | L* + a* + b* | 0.00400 | 0.00387 |

**Table S5.** The environmental parameters of the best regression model for estimated anthocyanin content and the effect of environmental factors in the model. 2x and 4x indicate diploid and allotetraploid, respectively. PL: precipitation lag, PW: precipitation window, TT: temperature threshold, TL: temperature lag, TW: temperature window, RL: radiation lag, RW: radiation window, P: precipitation, C: coldness (calculated from TT, TL, and TW), R: radiation, The unit is the number of days for all parameters. *: significant effect based on confidence intervals calculated with bias­ corrected and accelerated (BCa) bootstrapping.

| Site | Description | Genotype | Code | Best environmental parameter |  |  |  |  |  |  |  | Environmental factor effect |  |
| --- | --- | --- | --- | --- | --- | --- | --- | --- | --- | --- | --- | --- | --- |
|  |  |  |  | PL | PW | TT | TL | TW | RL | RW | P | C | R |
| Swiss | model plant 2x | *A. thaliana* | COL | 14 | 14 | 13 | 1 | 14 | 0 | 13 | * | N.S. | * |
| Swiss | progenitor 2x | *A. halleri* | HAL | 0 | 2 | 1 | 14 | 14 | 0 | 4 | N.S. | N.S. | * |
| Swiss | progenitor 2x | *A. lyrata* | MED | 1 | 1 | 13 | 1 | 14 | 0 | 1 | N.S. | * | * |
| Swiss | progenitor 2x | *A. lyrata* | SEP | 1 | 1 | 13 | 1 | 14 | 0 | 1 | N.S. | * | * |
| Swiss | synthetic 4x | *A. kamchatica* | RS7 | 0 | 1 | 1 | 14 | 14 | 0 | 2 | N.S. | * | * |
| Swiss | synthetic 4x | *A. kamchatica* | RS8 | 0 | 4 | 1 | 14 | 14 | 0 | 4 | * | * | * |
| Swiss | Northern 4x | *A. kamchatica* | DEN | 0 | 4 | 1 | 7 | 4 | 0 | 4 | * | * | * |
| Swiss | Northern 4x | *A. kamchatica* | POT | 0 | 1 | 1 | 14 | 14 | 0 | 2 | N.S. | * | * |
| Swiss | Japan 4x | *A. kamchatica* | MUR | 0 | 2 | 7 | 14 | 14 | 0 | 2 | N.S. | * | * |
| Swiss | Japan 4x | *A. kamchatica* | MAG | 0 | 6 | 1 | 14 | 14 | 0 | 6 | * | * | * |
| Swiss | Japan 4x | *A. kamchatica* | MAI | 1 | 3 | 13 | 2 | 14 | 0 | 4 | N.S. | * | * |
| Swiss | Japan 4x | *A. kamchatica* | TKS | 0 | 5 | 1 | 14 | 14 | 0 | 6 | N.S. | * | * |
| Japanese | model plant 2x | *A. thaliana* | COL | 2 | 2 | 1 | 0 | 2 | 0 | 4 | * | * | * |
| Japanese | progenitor 2x | *A. halleri* | HAL | 1 | 1 | 1 | 12 | 13 | 0 | 14 | N.S. | * | * |
| Japanese | progenitor 2x | *A. lyrata* | MED | 1 | 1 | 13 | 0 | 5 | 0 | 2 | N.S. | * | N.S. |
| Japanese | progenitor 2x | *A. lyrata* | SEP | 1 | 1 | 13 | 0 | 5 | 0 | 2 | * | * | N.S. |
| Japanese | synthetic 4x | *A. kamchatica* | RS7 | 0 | 4 | 4 | 5 | 5 | 0 | 4 | * | * | * |
| Japanese | synthetic 4x | *A. kamchatica* | RS8 | 1 | 1 | 4 | 0 | 14 | 0 | 14 | * | * | * |
| Japanese | Northern 4x | *A. kamchatica* | DEN | 1 | 1 | 13 | 0 | 8 | 0 | 2 | * | * | * |
| Japanese | Northern 4x | *A. kamchatica* | POT | 1 | 1 | 13 | 0 | 8 | 0 | 2 | N.S. | * | N.S. |
| Japanese | Japan 4x | *A. kamchatica* | MUR | 0 | 2 | 13 | 0 | 4 | 0 | 2 | * | * | * |
| Japanese | Japan 4x | *A. kamchatica* | MAG | 1 | 1 | 13 | 0 | 8 | 0 | 2 | N.S. | * | * |
| Japanese | Japan 4x | *A. kamchatica* | MAI | 1 | 1 | 13 | 0 | 14 | 0 | 2 | * | * | * |
| Japanese | Japan 4x | *A. kamchatica* | TKS | 0 | 2 | 13 | 0 | 5 | 0 | 2 | * | * | * |

**Table S6.** The settings of camera (RICOH WG-40) used for image acquisition. The Interval was set at either 90min Osee, or 60min Osee, depending on the routine at each site.

| Mode | Category | Subcategory | Setting |
| --- | --- | --- | --- |
| General shooting | General shooting mode |  | Interval shot |
|  | Setting of the Interval Shot | Interval | 90min 0sec |
|  |  | Number of shots | 1000 |
|  |  | Start Delay | Circumstance dependent |
|  | Fladh mode |  | Flash OFF |
| Recording | Image Tone |  | Natural |
|  | Recorded Pixel |  | 16M |
|  | Quality Level |  | ★★★ |
|  | White Balance |  | AWB |
|  | AF Setting | Focusing Area | whole |
|  |  | Auto Macro | ON |
|  |  | Focus Assist | OFF |
|  | AE Metering |  | Multi-segment metering |
|  | Sensitivity |  | AUTO |
|  | AUTO ISO Range |  | ISO125-1600 |
|  | EV Compensation |  | ±0.0 |
|  | D-Range Setting | Highlight | Auto |
|  |  | Shadow | Auto |
|  | Pixel Track SR |  | ON |
|  | Face Detection |  | OFF |
|  | Blink Detection |  | OFF |
|  | Digital Zoom |  | OFF |
|  | Instant Review |  | OFF |
|  | Memory | Make all items/settings | ON |
|  | Green Button |  | ON |
|  | Sharpness |  | 0 |
|  | Saturation |  | 0 |
|  | Contrast |  | 0 |
|  | Date Imprint |  | OFF |
|  | IQ Enhance |  | OFF |
|  | Macro Light |  | OFF |

**Table S7.**
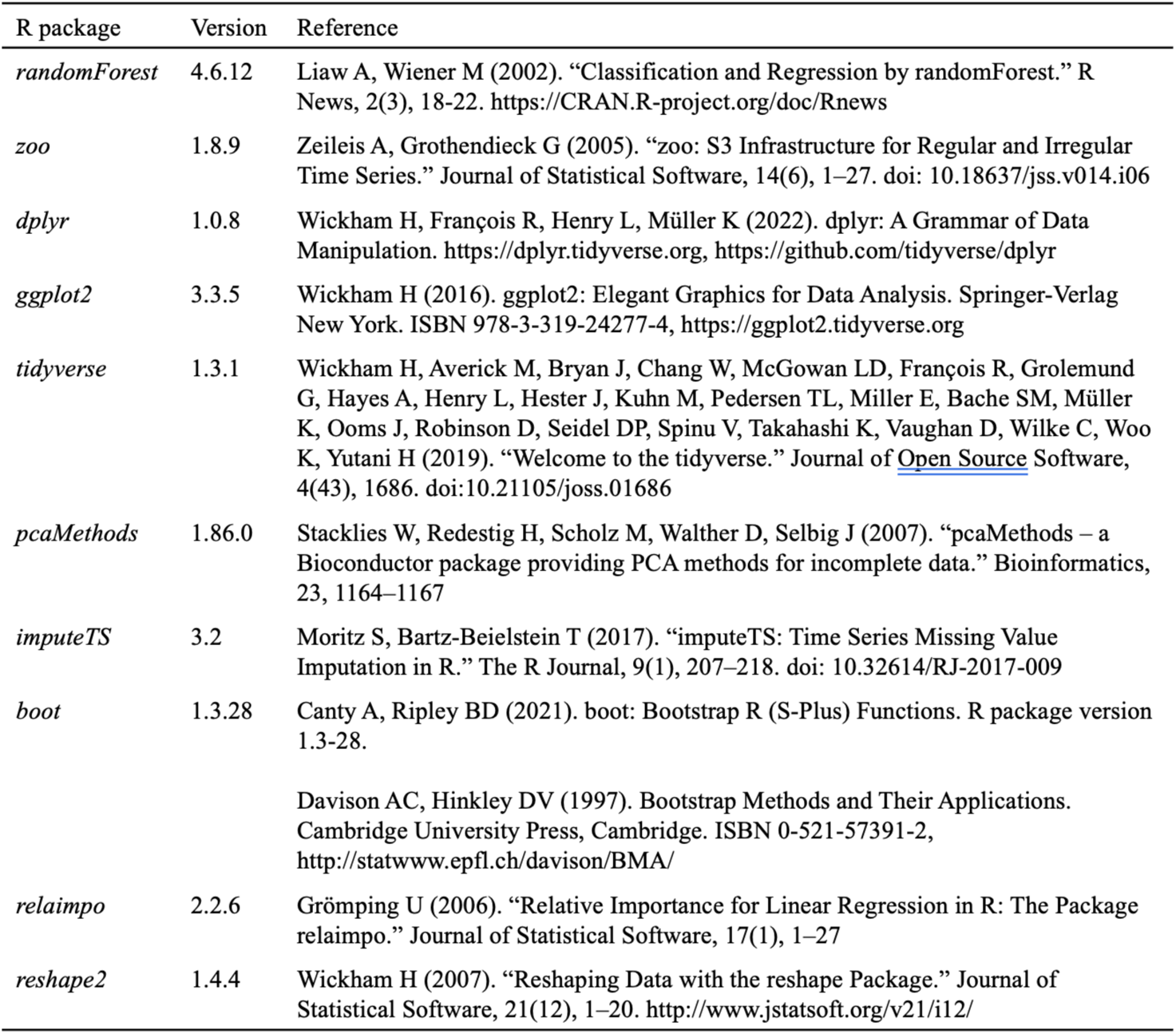
The details of the R packages used for data visualization and analyses.

**Table S8.** The summary of devices used for collecting environmental data at the Japanese site.

| Environmental factor | Product | Manufacturer | Distributor | Distributor location |
| --- | --- | --- | --- | --- |
| air temperature | HMP 155A | Campbell Scientific INC. | TAIYO KEIKI Co., Ltd. | Saitama |
| radiation | CMP-3 | PREDE Co., Ltd. | PREDE Co., Ltd. | Tokyo |
| precipitation | RT-5E | IKEDA KEIKI Co., Ltd. | IKEDA KEIKI Co., Ltd. | Tokyo |

## References

1. Kudoh, H. Molecular phenology in plants: In natura systems biology for the comprehensive understanding of seasonal responses under natural environments. New Phytol. 210, 399–412 (2016).

2. Suzuki, N., Rivero, R. M., Shulaev, V., Blumwald, E. & Mittler, R. Abiotic and biotic stress combinations. New Phytol. 203, 32–43 (2014).

3. Song, Y. H. et al. Molecular basis of flowering under natural long-day conditions in Arabidopsis. Nat. Plants 4, 824–835 (2018).

4. Shimizu, K. K., Kudoh, H. & Kobayashi, M. J. Plant sexual reproduction during climate change: Gene function in natura studied by ecological and evolutionary systems biology. Ann. Bot. 108, 777–787 (2011).

5. Yang, W. et al. Crop Phenomics and High-Throughput Phenotyping: Past Decades, Current Challenges, and Future Perspectives. Mol. Plant 13, 187–214 (2020).

6. Arya, S., Sandhu, K. S., Singh, J. & kumar, S. Deep learning: as the new frontier in high-throughput plant phenotyping. Euphytica 218, 1–22 (2022).

7. Weyler, J., Magistri, F., Seitz, P., Behley, J. & Stachniss, C. In-Field Phenotyping Based on Crop Leaf and Plant Instance Segmentation. Proc. - 2022 IEEE/CVF Winter Conf. Appl. Comput. Vision, WACV 2022 2968–2977 (2022) doi:10.1109/WACV51458.2022.00302.

8. Guo, W., et al. UAS-based plant phenotyping for research and breeding applications. Plant Phenomics 2021, (2021).

9. Virlet, N., Sabermanesh, K., Sadeghi-Tehran, P. & Hawkesford, M. J. Field Scanalyzer: An automated robotic field phenotyping platform for detailed crop monitoring. Funct. Plant Biol. 44, 143–153 (2017).

10. Hawkesford, M. J. & Lorence, A. Plant phenotyping: Increasing throughput and precision at multiple scales. Funct. Plant Biol. 44, v–vii (2017).

11. Egger, J. et al. Deep learning—a first meta-survey of selected reviews across scientific disciplines, their commonalities, challenges and research impact. PeerJ Comput. Sci. 7, 1–83 (2021).

12. Hüther, P., Schandry, N., Jandrasits, K., Bezrukov, I. & Becker, C. ARADEEPOPSIS, an automated workflow for top-view plant phenomics using semantic segmentation of leaf States. Plant Cell 32, 3674–3688 (2020).

13. Chang, S., Lee, U., Hong, M. J., Jo, Y. D. & Kim, J. B. Time-Series Growth Prediction Model Based on U-Net and Machine Learning in Arabidopsis. Front. Plant Sci. 12, 1–15 (2021).

14. Ronneberger, O., Fischer, P. & Brox, T. U-Net: Convolutional Networks for Biomedical Image Segmentation. in Medical Image Computing and Computer-Assisted Intervention (eds. Navab, N., Hornegger, J., Wells, W. & Frangi, A.) vol. 9351 (Springer, 2015).

15. Smith, A. G., Petersen, J., Selvan, R. & Rasmussen, C. R. Segmentation of roots in soil with U-Net. Plant Methods 16, 1–15 (2020).

16. Fan, D. P. et al. Camouflaged object detection. Proc. IEEE Comput. Soc. Conf. Comput. Vis. Pattern Recognit. 2774–2784 (2020) doi:10.1109/CVPR42600.2020.00285.

17. Mou, L. et al. CS2-Net: Deep learning segmentation of curvilinear structures in medical imaging. Med. Image Anal. 67, 101874 (2021).

18. Shakoor, N., Lee, S. & Mockler, T. C. High throughput phenotyping to accelerate crop breeding and monitoring of diseases in the field. Curr. Opin. Plant Biol. 38, 184–192 (2017).

19. Hrazdina, G. Anthocyanins. in The Flavonoids : Advances in Research (eds. Harborne, J. B. & Marby, T. J.) 135–186 (Chapman and Hall, 1982).

20. Chalker-Scott, L. Environmental significance of anthocyanins in plant stress responses. Photochem. Photobiol. 70, 1–9 (1999).

21. Manetas, Y. Why some leaves are anthocyanic and why most anthocyanic leaves are red? Flora Morphol. Distrib. Funct. Ecol. Plants 201, 163–177 (2006).

22. Nakabayashi, R. et al. Enhancement of oxidative and drought tolerance in Arabidopsis by overaccumulation of antioxidant flavonoids. Plant J. 77, 367–379 (2014).

23. Catalá, R., Medina, J. & Salinas, J. Integration of low temperature and light signaling during cold acclimation response in Arabidopsis. Proc. Natl. Acad. Sci. U. S. A. 108, 16475–16480 (2011).

24. Pescheck, F. & Bilger, W. High impact of seasonal temperature changes on acclimation of photoprotection and radiation-induced damage in field grown Arabidopsis thaliana. Plant Physiol. Biochem. 134, 129–136 (2019).

25. Van De Peer, Y., Mizrachi, E. & Marchal, K. The evolutionary significance of polyploidy. Nat. Rev. Genet. 18, 411–424 (2017).

26. Akagi, T., Jung, K., Masuda, K. & Shimizu, K. K. Polyploidy before and after domestication of crop species. Curr. Opin. Plant Biol. 69, 102255 (2022).

27. Shimizu, K. K. Robustness and generalist niche of polyploid species: genome shock or gradual evolution? Curr. Opin. Plant Biol. 69, 102292 (2022).

28. Gordon, S. P. et al. Gradual polyploid genome evolution revealed by pan-genomic analysis of Brachypodium hybridum and its diploid progenitors. Nat. Commun. 11, 1–16 (2020).

29. Burns, R. et al. Gradual evolution of allopolyploidy in Arabidopsis suecica. *Nat*. Ecol. Evol. 5, 1367–1381 (2021).

30. Stebbins, G. L. Chromosomal Evolution in Higher Plants. (1971).

31. Soltis, D. E., Visger, C. J. & Soltis, P. S. The polyploidy revolution then…and now: Stebbins revisited. Am. J. Bot. 101, 1057–1078 (2014).

32. Soltis, D. E., Visger, C. J., Blaine Marchant, D. & Soltis, P. S. Polyploidy: Pitfalls and paths to a paradigm. Am. J. Bot. 103, 1146–1166 (2016).

33. Shimizu, K. K., Fujii, S., Marhold, K., Watanabe, K. & Kudoh, H. Arabidopsis kamchatica (Fisch. ex DC.) K. Shimizu & Kudoh and A. kamchatica subsp. kawasakiana (Makino) K. Shimizu & Kudoh, New Combinations. Acta Phytotax. Geobot. 56, 163–172 (2005).

34. Hegarty, M. et al. Lessons from natural and artificial polyploids in higher plants. Cytogenet. Genome Res. 140, 204–225 (2013).

35. Hoffmann, M. H. Evolution of the realized climatic niche in the genus Arabidopsis (Brassicaceae). Evolution (N. Y). 59, 1425–1436. (2005).

36. Shimizu-Inatsugi, R. et al. The allopolyploid Arabidopsis kamchatica originated from multiple individuals of Arabidopsis lyrata and Arabidopsis halleri. Mol. Ecol. 18, 4024–4048 (2009).

37. Armstrong, J. J., Takebayashi, N. & Wolf, D. E. Cold tolerance in the genus Arabidopsis. Am. J. Bot. 107, 489–497 (2020).

38. Akama, S., Shimizu-Inatsugi, R., Shimizu, K. K. & Sese, J. Genome-wide quantification of homeolog expression ratio revealed nonstochastic gene regulation in synthetic allopolyploid Arabidopsis. Nucleic Acids Res. 42, (2014).

39. Paape, T. et al. Conserved but Attenuated Parental Gene Expression in Allopolyploids: Constitutive Zinc Hyperaccumulation in the Allotetraploid Arabidopsis kamchatica. Mol. Biol. Evol. 33, 2781–2800 (2016).

40. Paape, T. et al. Experimental and Field Data Support Range Expansion in an Allopolyploid Arabidopsis Owing to Parental Legacy of Heavy Metal Hyperaccumulation. Front. Genet. 11, 1–15 (2020).

41. Honjo, M. N. & Kudoh, H. Arabidopsis halleri: A perennial model system for studying population differentiation and local adaptation. AoB Plants 11, 1–13 (2019).

42. Kenta, T., Yamada, A. & Onda, Y. Clinal Variation in Flowering Time and Vernalisation Requirement across a 3000-M Altitudinal Range in Perennial Arabidopsis kamchatica Ssp. Kamchatica and Annual Lowland Subspecies Kawasakiana. J. Ecosyst. Ecography s6, 1–10 (2011).

43. Bomblies, K. & Madlung, A. Polyploidy in the Arabidopsis genus. Chromosom. Res. 22, 117–134 (2014).

44. Askey, B. C., Dai, R., Lee, W. S. & Kim, J. A noninvasive, machine learning–based method for monitoring anthocyanin accumulation in plants using digital color imaging. Appl. Plant Sci. 7, 1–8 (2019).

45. Chen, Y. Y. et al. Species-specific flowering cues among general flowering Shorea species at the Pasoh Research Forest, Malaysia. J. Ecol. 106, 586–598 (2018).

46. Saberioon, M., Amin, M. S. M., Gholizadeh, A. & Ezri, M. H. A review of optical methods for assessing nitrogen contents during rice growth. Appl. Eng. Agric. 30, 657–669 (2014).

47. Livne, M. et al. A U-net deep learning framework for high performance vessel segmentation in patients with cerebrovascular disease. Front. Neurosci. 13, 1–13 (2019).

48. Bardis, M. et al. Deep learning with limited data: Organ segmentation performance by U-net. Electron. 9, 1–12 (2020).

49. Samarasinghe, G. et al. Deep learning for segmentation in radiation therapy planning: a review. J. Med. Imaging Radiat. Oncol. 65, 578–595 (2021).

50. Liu, J. & Wang, X. Plant diseases and pests detection based on deep learning: a review. Plant Methods 17, 1–18 (2021).

51. Fu, J. et al. Dual attention network for scene segmentation. Proc. IEEE Comput. Soc. Conf. Comput. Vis. Pattern Recognit. 2019-June, 3141–3149 (2019).

52. Garcia-Garcia, A. et al. A survey on deep learning techniques for image and video semantic segmentation. Appl. Soft Comput. J. 70, 41–65 (2018).

53. Zheng, X. T. et al. The major photoprotective role of anthocyanins in leaves of Arabidopsis thaliana under long-term high light treatment: antioxidant or light attenuator? Photosynth. Res. 149, 25–40 (2021).

54. Aikawa, S., Kobayashi, M. J., Satake, A., Shimizu, K. K. & Kudoh, H. Robust control of the seasonal expression of the Arabidopsis FLC gene in a fluctuating environment. Proc. Natl. Acad. Sci. U. S. A. 107, 11632–11637 (2010).

55. Nagano, A. J. et al. Annual transcriptome dynamics in natural environments reveals plant seasonal adaptation. Nat. Plants 5, 74–83 (2019).

56. Nishio, H. et al. Repressive chromatin modification underpins the long-term expression trend of a perennial flowering gene in nature. Nat. Commun. 11, (2020).

57. Yamaguchi, N. et al. H3K27me3 demethylases alter HSP22 and HSP17.6C expression in response to recurring heat in Arabidopsis. Nat. Commun. 12, 1–16 (2021).

58. Paape, T. et al. Patterns of polymorphism and selection in the subgenomes of the allopolyploid Arabidopsis kamchatica. Nat. Commun. 9, (2018).

59. Takahagi, K. et al. Homoeolog-specific activation of genes for heat acclimation in the allopolyploid grass Brachypodium hybridum. Gigascience 7, 1–13 (2018).

60. Sun, J. et al. A Recently Formed Triploid Cardamine insueta Inherits Leaf Vivipary and Submergence Tolerance Traits of Parents. Front. Genet. 11, 1–12 (2020).

61. An, N. et al. Plant high-throughput phenotyping using photogrammetry and imaging techniques to measure leaf length and rosette area. Comput. Electron. Agric. 127, 376–394 (2016).

62. Stockenhuber, R. et al. The UV RESISTANCE LOCUS 8-mediated UV-B response is required alongside CRYPTOCHROME1 for plant survival under sunlight in the field. bioRxiv 1–36 (2021).

63. Naik, H. S. et al. A real-time phenotyping framework using machine learning for plant stress severity rating in soybean. Plant Methods 13, 1–12 (2017).

64. Ebersbach, J. et al. Exploiting High-Throughput Indoor Phenotyping to Characterize the Founders of a Structured B. napus Breeding Population. Front. Plant Sci. 12, (2022).

65. Wang, C. et al. A cucumber leaf disease severity classification method based on the fusion of DeepLabV3+ and U-Net. Comput. Electron. Agric. 189, 106373 (2021).

66. Briskine, R. V. et al. Genome assembly and annotation of Arabidopsis halleri, a model for heavy metal hyperaccumulation and evolutionary ecology. Mol. Ecol. Resour. 17, 1025–1036 (2017).

67. Shimizu-Inatsugi, R. et al. Metal accumulation and its effect on leaf herbivory in an allopolyploid species Arabidopsis kamchatica inherited from a diploid hyperaccumulator A. halleri. Plant Species Biol. 36, 208–217 (2021).

68. Cui, Z., Yang, J. & Qiao, Y. Brain MRI segmentation with patch-based CNN approach. in Chinese Control Conference, CCC vols 2016-Augus 7026–7031 (TCCT, 2016).

69. He, K., Zhang, X., Ren, S. & Sun, J. Deep residual learning for image recognition. Proc. IEEE Comput. Soc. Conf. Comput. Vis. Pattern Recognit. 2016-Decem, 770–778 (2016).

70. Tan, M. & Le, Q. V. EfficientNet: Rethinking model scaling for convolutional neural networks. 36th Int. Conf. Mach. Learn. ICML 2019 2019-June, 10691–10700 (2019).

71. Schmidt, R. & Mohr, H. Time-dependent changes in the responsiveness to light of phytochrome-mediated anthocyanin synthesis. Plant, Cell Environ. Cell Environ. 4, 433–437 (1981).

72. Breiman, L. Random Forests. Mach. Learn. 45, 5–32 (2001).

73. Ali, M., Borgo, R. & Jones, M. W. Concurrent time-series selections using deep learning and dimension reduction. Knowledge-Based Syst. 233, 107507 (2021).

74. Mohtashemi, M., Kleinman, K. & Yih, W. Multi-syndrome analysis of time series using PCA: A new concept for outbreak investigation. Stat. Med. 26, 5203–5244 (2007).

75. Black, B. A. et al. Winter and summer upwelling modes and their biological importance in the California Current Ecosystem. Glob. Chang. Biol. 17, 2536–2545 (2011).

76. Catalá, R., Medina, J. & Salinas, J. Integration of low temperature and light signaling during cold acclimation response in Arabidopsis. Proc. Natl. Acad. Sci. U. S. A. 108, 16475–16480 (2011).

77. Olsen, K. M., Lea, U. S., Slimestad, R., Verheul, M. & Lillo, C. Differential expression of four Arabidopsis PAL genes; PAL1 and PAL2 have functional specialization in abiotic environmental-triggered flavonoid synthesis. J. Plant Physiol. 165, 1491–1499 (2008).

78. Petridis, A., Döll, S., Nichelmann, L., Bilger, W. & Mock, H. P. Arabidopsis thaliana G2-LIKE FLAVONOID REGULATOR and BRASSINOSTEROID ENHANCED EXPRESSION1 are low-temperature regulators of flavonoid accumulation. New Phytol. 211, 912–925 (2016).

79. Havaux, M. & Kloppstech, K. The protective functions of carotenoid and flavonoid pigments against excess visible radiation at chilling temperature investigated in Arabidopsis npq and tt mutants. Planta 213, 953–966 (2001).

80. Zhang, Y., Zheng, S., Liu, Z., Wang, L. & Bi, Y. Both HY5 and HYH are necessary regulators for low temperature-induced anthocyanin accumulation in Arabidopsis seedlings. J. Plant Physiol. 168, 367–374 (2011).

81. Schmidt, R. & Mohr, H. Time-dependent changes in the responsiveness to light of phytochrome-mediated anthocyanin synthesis. Plant. Cell Environ. 4, 433–437 (1981).

## References

1. Schmidt R, Mohr H. Time-dependent changes in the responsiveness to light of phytochrome-mediated anthocyanin synthesis. Plant Cell Environ. 1981;4(6):433–7.

2. Schindelin J, Arganda-Carreras I, Frise E, Kaying V, Longair M, Pietzsch, T, Preibisch S, Rueden C, Saalfeld S, Schmid B, Tinevez JY, White DJ, Hartenstein V, Eliceiri K, Tomancak P, Cardona A. Fiji: An open-source platform for biological-image analysis. Nature Methods. 2012; 5(7):676–682.

